# Acute E2/P4 loss compromises the biology and function of neurogenic niches during a vulnerable female aging period

**DOI:** 10.1101/2025.05.06.652460

**Authors:** Sree Vani Pillutla, Maddy J Skoda, Atsushi Ishii, Mandi J Corenblum, Tyler Montogomery, Paige Wene, Nainika Menakuru, Jennifer N Kraszewski, Dean Billheimer, Gabriel M Winter, Stephen L Cowen, Lalitha Madhavan

## Abstract

Effects of aging on neural stem progenitor cells (NSPCs) have been studied in males, but less is known in females. Here we comparatively assess female NSPC biology, both in the subventricular zone and hippocampal dentate gyrus niches, across different ages of F344 rats (2, 6, 9 and 14 months). The rats were ovariectomized (OVX) or remained Intact at each of the aging stages, to assess the role of the female sex hormones, estradiol (E2) and progesterone (P4). Results show that while age-dependent decays become prominent at 14 months, ovariectomy-induced E2/P4 loss markedly reduces neurogenesis and associated behavioral function, earlier, at 9 months of age. Coinciding with this pattern of neurogenic decline, we also detect adaptive changes in estrogen and progesterone receptor expression, antioxidant expression, and brain E2/P4 levels. Fundamentally, these results reveal specific female time-periods, when the brain is sensitive to age and E2/P4 loss, potentially ‘setting-up’ for disease susceptibility.

## INTRODUCTION

The adult rodent brain contains actively dividing neural stem progenitor cells (NSPCs) that retain a lifelong capacity for neurogenesis (Alvarez-Buylla and Lim, 2004; Bond et al., 2015). This is a core plastic process proposed to be a neural reserve that can be called upon at times of high physiological demand or pathology (Kempermann, 2008; 2015). Broadly, age and sex dependent processes are known to modulate the behavior and function of adult NSPCs (Katsimpardi and Lledo, 2018; Liu and Rando, 2011; Yagi and Galea, 2019; Zhao et al., 2021). However, these influences are not fully understood. It is recognized that neurogenesis declines with age in both the major NSPC niches – the forebrain subventricular zone (SVZ) and the subgranular zone (SGZ) of the hippocampal dentate gyrus (DG) – and there has been much scrutiny on the mechanisms driving these decreases. In this context, our previous work identified a critical period of decline in NSPC regenerative function in male rodents and discovered the reduced expression of the nuclear factor (erythroid-derived 2)-like 2 (Nrf2) transcription factor as a key mechanism driving this process (Anandhan et al., 2021a; Corenblum et al., 2016; Dodson et al., 2021; Ray et al., 2018).

However, although the effects of age on neurogenesis in males have been examined, female NSPC aging remains underexplored. Thus far, studies point to sex hormones and their receptors as important differential regulators of adult neurogenesis in females (Mahmoud et al., 2016; Ponti et al., 2018). Circulating hormone levels of 17b-estradiol (E2) and progesterone (P4) fluctuate dramatically during a female rodent’s life during breeding, estrous cycle, pregnancy, lactation, and menopause (Pawluski et al., 2009). Moreover, species and strain differences further complicate these hormonal stages. Despite these complexities, E2 and P4 have been studied in relation to some of these natural variations and they have been shown enhance NSPC function in most cases. Generally, NSPC proliferation has been reported to be the greatest during the proestrus phase of the estrous cycle, when E2 levels are particularly high, with P4 also appearing to enhance NSPC activity (Ponti *et al*., 2018; Zhang et al., 2010). Sex-hormone levels, however, change across the lifespan and the question of how age modulates female NSPCs has not been carefully addressed.

It is also not understood whether female NSPC biology differs between the two adult NSPC niches in the SVZ and hippocampal DG. Elegant studies, mostly focused on hippocampal NSPCs, have reported basal sex differences in cell proliferation but not in survival of new neurons in the DG (O’Leary et al., 2022; Seib et al., 2018; Yagi et al., 2016; Yagi et al., 2020). For example, it has been reported that adult female rats have ∼50% more newly proliferating cells and fewer dying cells in the DG during proestrus (high estradiol levels) compared to males or adult females in estrus or diestrus stages (low estradiol levels) (Pawluski *et al*., 2009). However, SVZ NSPC dynamics in females has not been well characterized and what happens with age, comparatively between the SVZ and DG niches, has not been assessed.

To address these questions, the current study investigated the age-related changes in neurogenesis, and its related physiology and behavior, in different age-groups of female rats - F344 rats aged 2, 6, 9 and 14 months (mos). We also assessed the influence of female sex hormones at the different ages by inducing an acute loss of E2 and P4 via ovariectomy (a model of surgical menopause). About two weeks after ovariectomy, changes in both the SVZ and DG NSPC niches were compared. Results show that although neurogenesis broadly decreases with age, a vulnerable phase emerges around 9 months of age when E2/P4 loss compromises neurogenesis and related behavioral function. The study also establishes parallel changes in E2/P4 receptor expression and antioxidant expression and also examines estrous cycle and brain E2/P4 concentration changes, in the four age-groups. These results, for the first time, provide a comparative profile of both the SVZ and DG neurogenic niches in the context of aging and sex hormone influences.

## RESULTS

### Ovariectomy compromises NSPC-relevant behavioral function in aging female rats

We assessed the behavioral and pathological effects of acute E2 and P4 loss, utilizing intact versus ovariectomized rats aged 2, 6, 9, and 14 months (mos) of age (**Fig. 1A**). Female F344 rats underwent bilateral ovariectomy (OVX) or sham surgery (Intact) and were assessed 2 weeks later through NSPC specific behavioral tasks, namely fine olfactory discrimination (FOD; SVZ correlate), object-context pattern separation (OCPS, DG correlate), and reversal learning in the Morris water maze (RMWM, DG correlate) (LM refs). Subsequently, rat tissues were subjected to different cellular and molecular assays including estrous cycle staging and measurement of brain E2 and P4 levels.

**Figure 1.**
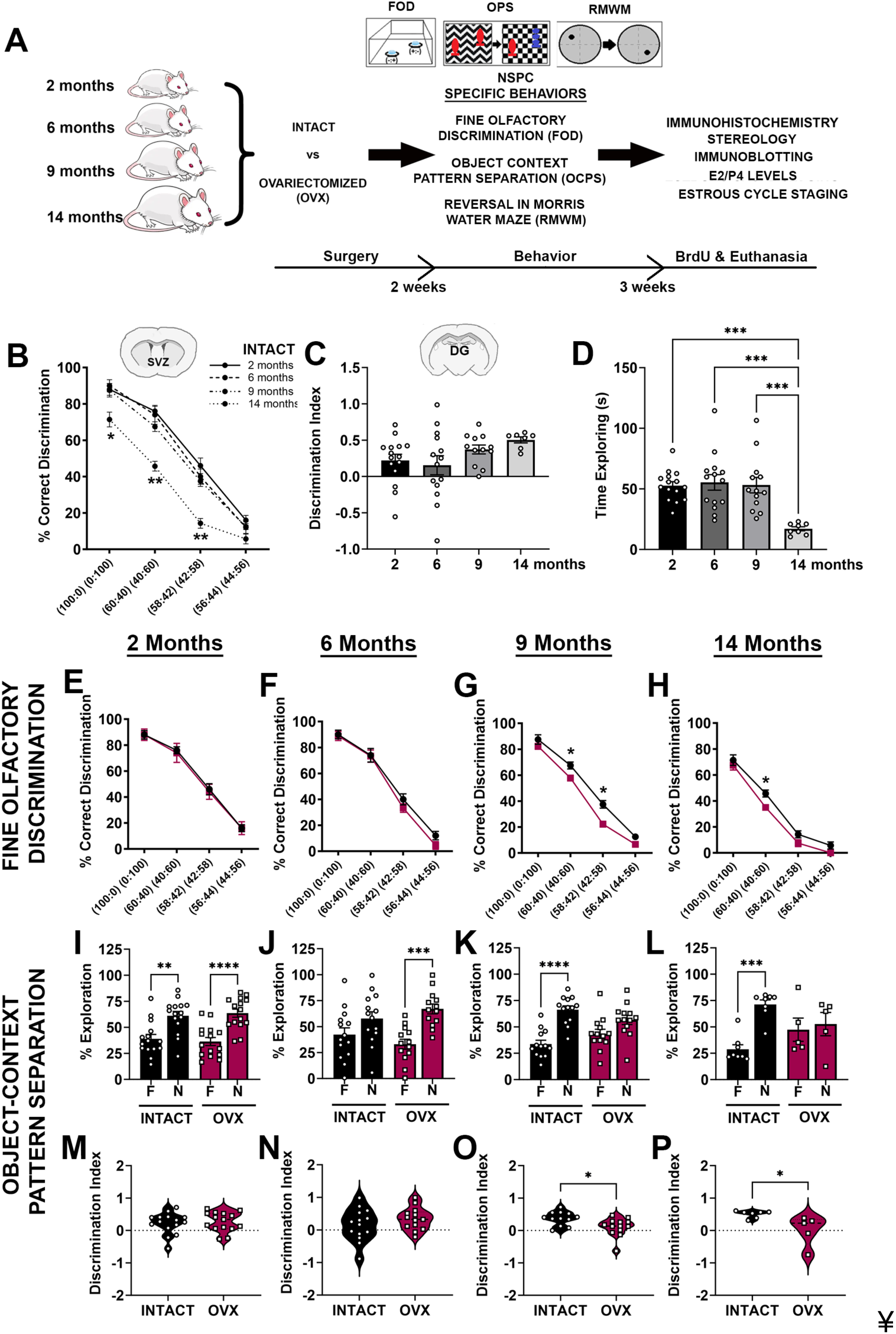
Effects of age and ovariectomy on NSPC relevant behaviors. The schematic in (A) outlines the overall experimental design of the study and the specific outcome assays performed. (B) shows results across the four age-groups (2, 6, 9, 14 mos) of Intact female F344 rats assessed through FOD task. The results from the OCPS task, with respect to age are in (C) and (D) in terms of the Discrimination Index (DI) and the time spent exploring the objects. Comparison of FOD performance between Intact and OVX rats at the 4 different ages is in (E-H). Similarly, a comparison of the OCPS behavioral performance between Intact and OVX at the four ages is in (I-P). OCPS results are expressed as a DI (I-L, DI) and the percentage of time spent exploring the novel versus familiar object (M-P). **** p < 0.0001, *** p = 0.001, ** p < 0.01, * p < 0.05; One-way ANOVA with Tukey’s post-hoc test (C, D), Two-way ANOVA with Tukey’s post-hoc test (B, E-L), Unpaired t-tests (M-P); mean ± SEM, n = 7-10 animals/group in FOD, n = 7-15 animals/group in OCPS.

We first assessed the rat’s ability to discriminate between different ratios of [+]/good tasting coconut (COC) and [−]/bad tasting mixture of almond and denatonium benzoate (ALM), via the FOD task. Comparing the percent of correct discriminations across the age groups in Intact animals, and a prominent reduction was seen at 14 mos. The 14 mos old rats performed significantly worse than their younger counterparts at both discrete (100:0) and fine discrimination (60:40 and 58:42) concentrations (**Fig. 1B**). Specifically, at the 100:0 odorant concentration, 14 mos old animals correctly discriminated ∼71% of the time versus ∼90% in 6 mos old animals (p = 0.0322, F_3,124_ = 334.4 (concentration), two-way ANOVA with Tukey’s multiple comparisons). Compared to their 9 mos old littermates, FOD was compromised in the 14 mos old rats at both the 60:40 (67.5% and 45.71%) and 58:42 (37.5% and 14.29%) odorant ratios (p = 0.0078 and p = 0.0002, two-way ANOVA with Tukey’s multiple comparisons). In the context of acute hormone loss, no significant alterations upon ovariectomy were noted between Intact and OVX rats at 2 and 6 mos of age (**Fig. 1E, F**). However, 9 mos old OVX rats showed significant deficits in their ability to discriminate between very similar ratios of COC and ALM (60:40 and 58:42) compared to their Intact counterparts (**Fig. 1G**; 60:40 ratio – p=0.034, t=2.331, df=15; 58:42 ratio – p=0.022, t=2.537, df=15, Unpaired t-tests). FOD was also stunted in ovariectomized animals at 14 months of age at the 60:40 odor ratio (**Fig. 1H**; p=0.048, t=2.181, df=13, Unpaired t-test).

DG NSPC function was tested through an object-context pattern separation (OCPS) task. Specifically, we measured the animal’s ability to differentiate specific stimuli (sprocket tower vs figurine statue) within highly similar environments (chevron vs concentric circle floor patterns). Animals with durable hippocampal neurogenesis would be expected to spend more time exploring the novel object (the stimuli ultimately paired up with the novel floor pattern). Across age, Intact animals showed no differences in the Discrimination Index (DI, **Fig. 1C**), however the total time spent by the rats exploring the objects decreased significantly at 14 months of age (**Fig. 1D**; p = 0.0005, F_3,46_ = 8.631, one-way ANOVA with Tukey’s post-hoc test). DI is a subsequent measure that corrects for the total exploration time. A higher DI indicates better differentiation between the target object (stimuli seen during encoding but in a different context) and the familiar object being presented.

When the percent of time exploring the novel vs familiar object was assessed, all Intact groups explored the novel object much more than the familiar object (**Fig. 1I-L**). However, in the OVX groups, although significantly more time was spent with the novel object at ages 2 and 6 months (**Fig 1I** - p = 0.001, F_3,50_ = 6.699; **Fig 1J** - p = 0.001, F_3,56_ = 12.52, one-way ANOVA with Tukey’s post-hoc test), no significant differences were found between novel and familiar object explorations at 9 and 14 mos of age (**Fig. 1K, L)**. With respect to the DI, at 2 and 6 months, no differences were noted between Intact and OVX groups (**Fig. 1M-N**; p = 0.6521 and p = 0.2713 respectively, Unpaired t-tests). But, at 9 months, OVX rats showed a significantly lower DI than the Intact rats (**Fig. 1O**; p = 0.0147, t=2.639, df=23, Unpaired t-test). Similarly, the OVX animals at 14 months of age had a diminished discrimination index when compared to Intact (**Fig. 1P**; p = 0.0204, t=2.753, df=10, Unpaired t-test). These data suggest that by 9 months of age, female rats with an acute loss of circulating sex hormones, demonstrate substantial declines in both SVZ and DG NSPC-related behavioral function.

### Ovariectomy affects spatial learning and alters the search strategies employed by the ovariectomized rats especially at 9 months of age

The Morris water maze with a reversal learning aspect was implemented to examine spatial learning and cognitive flexibility in the rats. We and others have reported that animals with suppressed adult hippocampal neurogenesis show impairments in relearning a new goal position after platform reversal in the Morris water maze (Garthe et al., 2014; Garthe and Kempermann, 2013; Ray *et al*., 2018). ANY-maze software (Stoelting Co., Wood Dale, IL, USA) was used to run trials and calculate the corrected integrated path length (CIPL), to the hidden platform, which corrects for swim speed and release location. Generally, an age-related decline in both the acquisition (first 4 days) and reversal learning (days 5 and 6) were noted in the Intact animals (**Fig. 2A**, p=0.044, F_15,318_ = 2.284 (interaction), two-way ANOVA). When the Intact rats were compared to their OVX counterparts at the different ages, in terms of the initial acquisition phase (first 4 days), the 9 mos old OVX rats showed higher CIPL scores than the Intact animals (day 3: two-way RM-ANOVA with Šídák’s multiple comparisons test, p=0.0287, t=2.862, df=144), while no significant differences were seen at the other ages (**Fig. 2C-F**; two-way RM-ANOVA, group X time is significant at p=0.0153). Upon goal reversal (day 5), the 9 mos old Intact rats showed significantly higher CIPL scores than the OVX animals (two-way RM-ANOVA with Šídák’s multiple comparisons test, p=0.0211, t=2.965, df=144). On the other hand, 14 mos old Intact animals showed the opposite trend and had lower CIPL scores on day 5 than their OVX counterparts (p=0.0352, t=2.789, df=156, two-way RM-ANOVA with Šídák’s multiple comparisons test).

**Figure 2.**
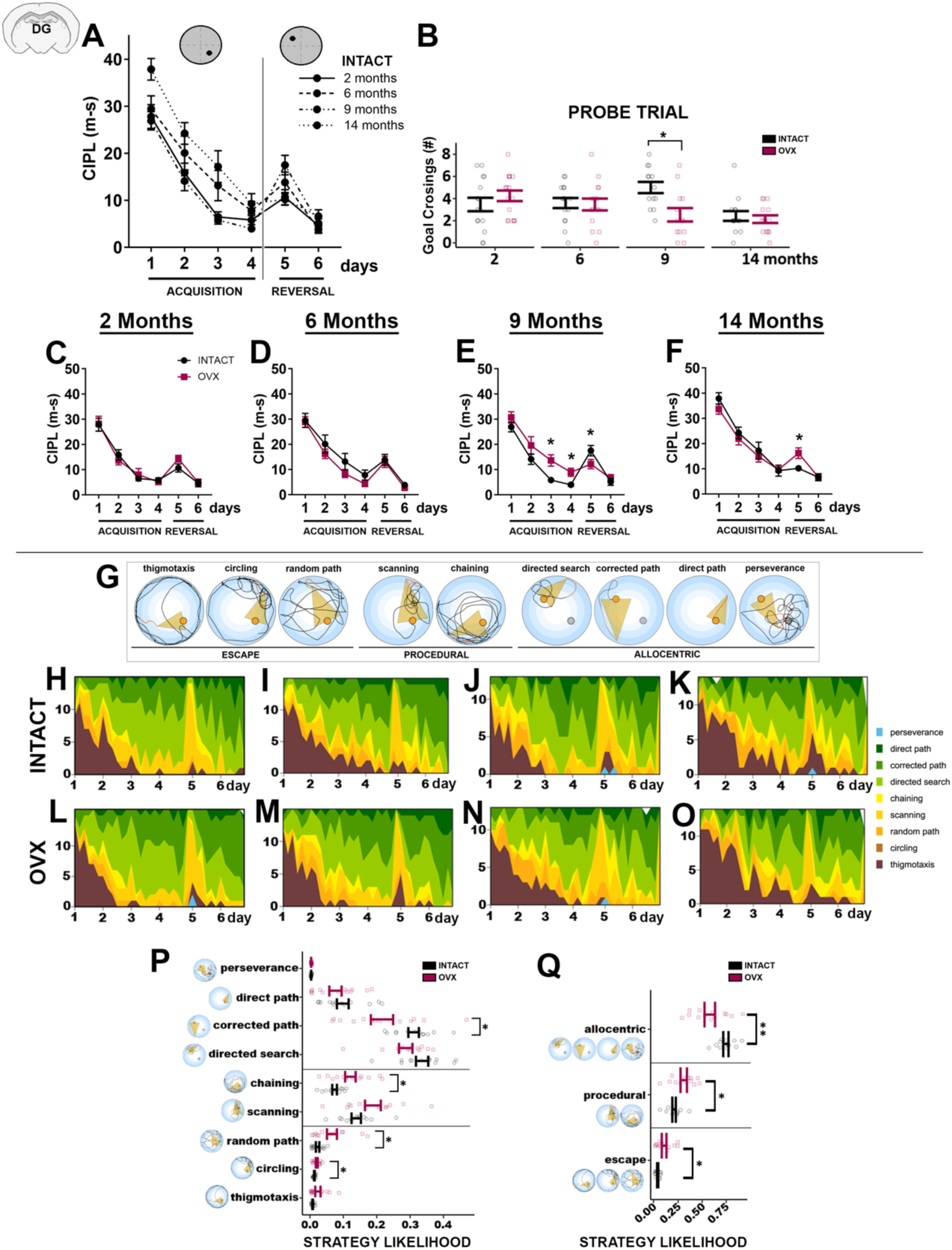
Influences of aging and E2/P4 loss on spatial learning and strategy usage. (A) compares the performance of the 2, 6, 9 and 14 mos old Intact female rats in the RMWM task which assesses spatial learning and cognitive flexibility. The results of the probe trial conducted after the acquisition phase, at the end of day 4, is shown in (B). The differences in RMWM performance between Intact and OVX rats at the four age-groups, across the acquisition and platform reversal phases, is displayed in (C-F). (G) outlines the 9 different swimming strategies employed by the rats that were analyzed. The strategy analyses plots comparing Intact vs OVX groups at 2, 6, 9 and 14 mos of age are in (H-O). Line separates Allocentric, Procedural and Escape strategies. ** p < 0.01, * p < 0.05; Two-way ANOVA with Tukey’s post-hoc test (A), Two-way RM-ANOVA with Šídák’s multiple comparisons test (C-F); Two-way ANOVAs and Unpaired t-tests (B, P, Q); mean ± SEM, n = 13-16 animals/group.

To further assess acquisition performance, goal crossings were analyzed during the probe trial administered on day 4. While no differences between OVX and Intact animals were observed on months 2, 6, and 14, goal crossings were significantly reduced in the 9 mos OVX group, indicating that these animals had not learned the location of the hidden platform well (**Fig. 2B**, p=0.019, t=3.12, df = 23.27, Holm-Bonferroni correction for 4 comparisons).

The mean allocentric confidence on day 5 was also significantly lower in the 9 mos OVX group compared to Intact in the probe trial at the end of day 4 (**Supp. Fig. 1A**). No differences between 9 and 14 mos old rats were found in the probe trial at the end of day 6 (**Supp. Fig. 1B**). More broadly, when the correlation between CIPL scores and the probability of using an allocentric strategy was plotted, an expected strong and negative relationship between CIPL scores and the likelihood of selecting an allocentric strategy was seen (**Supp. Fig. 1C**).

To develop a more fine-grained understanding of the behavioral strategy employed by each animal, we measured the likelihood (confidence) the animal was using a specific search strategy to find the platform using the algorithm described in Garthe et al., [(Garthe et al., 2009), also see Methods]. This approach allows the identification of 9 unique search strategies during water maze performance. Each of these strategies falls into the broader categories of non-goal-directed, procedural, and allocentric search, with the allocentric strategy being the most effective for finding the platform (see **Schemas in Fig. 2G)**. **Figure 2H-O** plots each group’s strategy usage across all trials over the 6 days (excluding probe trials), showing a shift in all groups from non-goal directed strategies to allocentric strategies (green). We also found that the mean likelihood OVX animals were using an allocentric strategy on day 5 was significantly lower in the 9 mos animals when compared to Intact in the probe trial at the end of day 4 (**Supp. Fig. 1A**). No differences between 9 and 14 mos old rats were found in the probe trial at the end of day 6 (**Supp. Fig. 1B**). We also observed that the use of the allocentric strategy for animals in the OVX and Intact groups was, as expected, negatively correlated with the CIPL scores (**Supp. Fig. 1C**).

Overall, both groups exhibited a gradual shift towards allocentric strategies over the 4-day acquisition phase, and between-group differences were observed in 9 mos old animals. Specifically, on day 4, the 9 mos old OVX rats used significantly less corrected path strategy (p = 0.01) and more chaining (p = 0.01), random path (p = 0.01) and circling strategies (p = 0.02) (**Fig. 2P**) compared to Intact rats. These data broadly translated to lesser usage of allocentric strategies (p = 0.01) and more procedural (p = 0.02) and escape strategies (p = 0.01) by the 9 mos old OVX rats (**Fig. 2Q**, two-way ANOVA with Holm-Sidak multiple comparisons test).

Using Rtrack (see Methods) we generated density plots to visualize the proportion of time the rats spent at each location in the pool. Density plots from a subset of animals in the OVX and Intact groups are presented in **Figure 3A-D**. The figure shows a composite of all trials for each animal on Days 4 and 5 with orange indicating a higher proportion of time spent by the animals at a given location. In the 9 mos old rats, we found a significance difference by group (OVX vs. Intact) with respect to the percentage of time spent in the goal quadrant (**Figure 3E**, two-way RM-ANOVA, F(1) = 14.25, p = 0.00093, η^2^ = 0.37). We also assessed the likelihood the animals were using an allocentric strategy (Allocentric Confidence) by adding the likelihoods animals were using the directed path, directed search, and corrected path strategies. Here again, the 9 mos old animals rats showed a clear difference in that OVX group had significantly less allocentric confidence than the Intact animals (two-way RM-ANOVA, F(1) = 6.37, p = 0.01, η^2^ = 0.21). Overall, these data show that, particularly at 9 mos of age, intact rats used more allocentric strategies to find the location of the platform and learn the task better than the ovariectomized animals.

**Figure 3.**
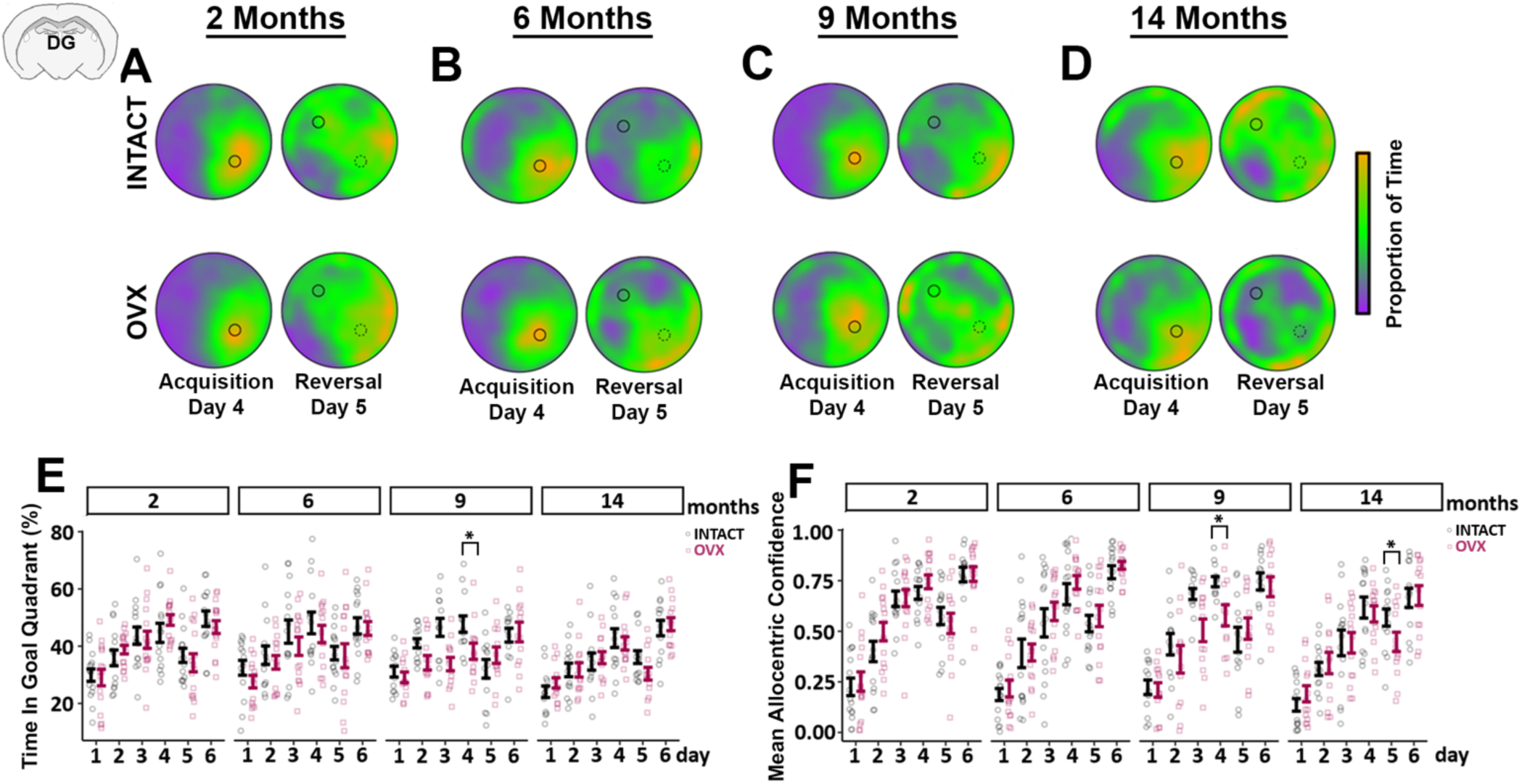
Ovariectomy-induced differences in goal quadrant time and allocentric strategy use in the aging female rats. (A-D) exhibits example density plots from 2, 6, 9 and 14 mos old female rats that indicates the proportion of time spent in the different areas of the pool on day 4 and day 5 of the RMWM task. Orange indicates a higher proportion of time spent in an area whereas purple indicates a lower proportion. (E) shows a comparison of time spent in the goal quadrant on each of the 6 days in the four aging stages. The mean allocentric confidence results are shown in (F). ** p < 0.01, * p < 0.05; Two-way ANOVAs and Unpaired t-tests (E, F); mean ± SEM, n = 13-16 animals/group.

### Age effects on estrous cycle states and brain E2/P4 levels and their relationship to neurogenesis-relevant behaviors

We examined estrous cycle stages through vaginal cytology, and measured E2 and P4 levels in brain tissues via LC-MS/MS, in the four age groups of Intact rats (**Fig. 4**). For estrous cycle staging vaginal samples were collected at multiple times as described in the methods and the number of animals in each age/group at the different estrous cycle stages was computed (**Fig. 4A**). As shown in Figures **4B-E**, 2 and 6 mos old animals were found consistently in all the four phases of proestrus, estrus, metestrus, and diestrus across time suggesting that they were regularly cycling. In contrast, many of the older animals (9 and 14 mos) were found in diestrus with a few in estrus (**Fig. 4B-E**). Some 9 mos old rats continued to have regular cycles. Most of the 14 mos old rats appeared to have entered an extended estrus phase, besides a few that continued to have regular cycles. No significant relationships were found between estrous stages and task performance in the FOD (**Fig. 4F**, analysis at the finer 58:42 and 56:44 odor ratios) and discrimination index (DI) in the OCPS (**Fig 4G**). ANOVA-based linear regression analysis showed that age significantly associated with olfactory discrimination (OD) at the 58:42 ratio (p = 0.003) in the full model and also in the reduced model (p = 0.0004). However, estrous stage did not significantly predict OD at either concentration. ANOVA results from linear regression to determine effects of estrous stage and age for the OCPS task indicated that neither age nor estrous stage were significant predictors of DI.

**Figure 4:**
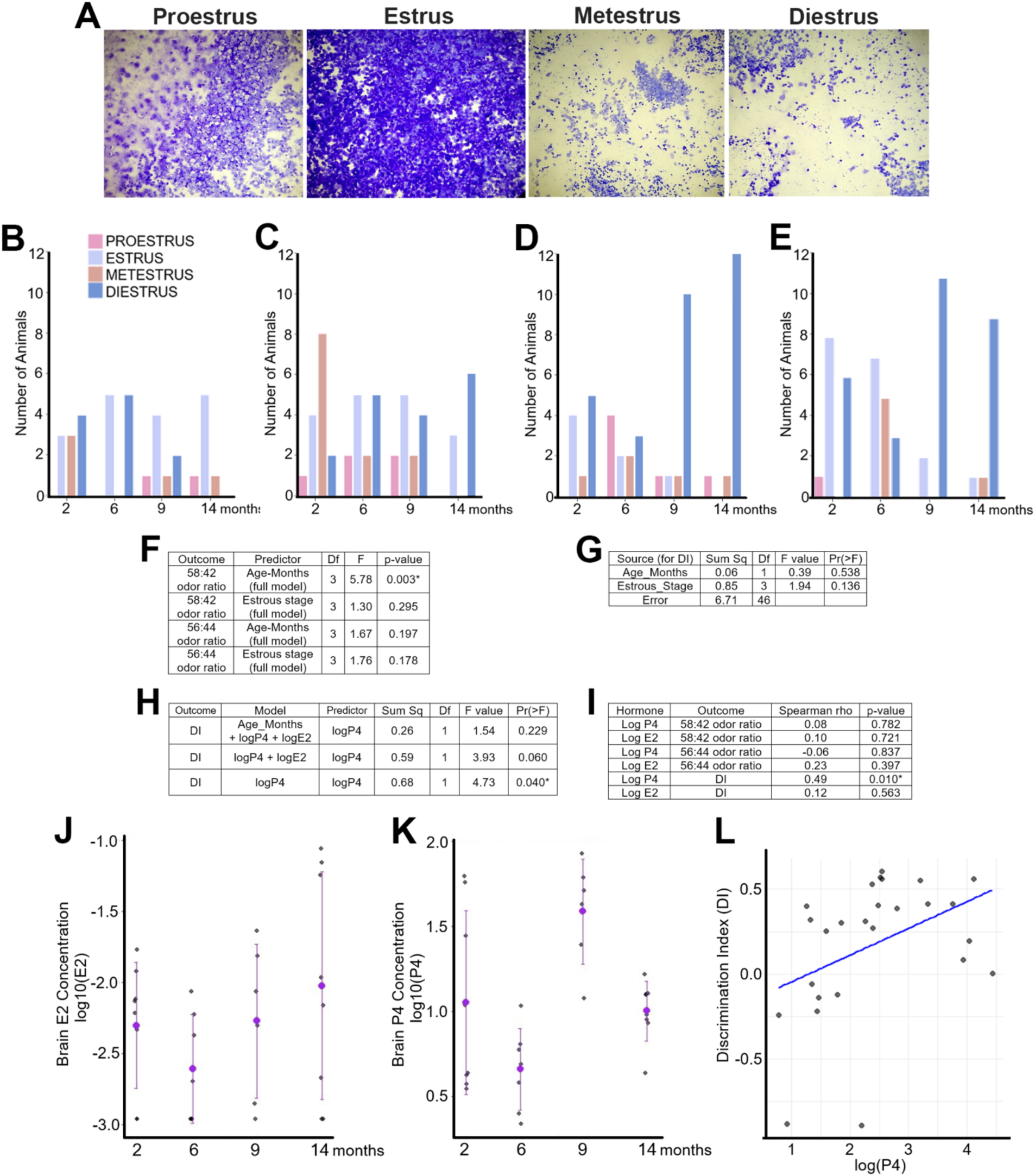
Characterization of estrous cycle stages and E2/P4 levels with age and their relationship with behavior. (A) depicts vaginal cytology defining the 4 estrous cycle stages. (B-E) characterize the number of Intact animals in the different estrous stage by age at 4 timepoints (after FOD, OCPS, RMWM and before euthanasia). (F-G) show how brain levels of E2 and P4 vary across the 4 age-groups in Intact rats. These are log10-transformed E2 and P4 concentrations by age where the purple point is the mean value, and the error bars are based on standard deviation (SD). The relationship between P4 concentrations and discrimination index in OCPS is shown in (H). Summary of ANOVA from linear models using olfactory discrimination of 58:42 and 56:44 tasks is in (I), and (J) shows ANOVA results from linear regression to determine effects of estrous stage and age on DI for the pattern separation task. ANOVA results from linear regression to determine effects of P4, E2 and age on DI for the pattern separation task are in (K). The table in (L) conveys the Spearman’s rank correlation to assess the relationship between brain E2 and P4 levels and the outcomes in FOD (2 ratios of 58:42 and 56:44) and OCPS (DI).

LC-MS/MS based measurement of E2/P4 levels showed high variability between animals in each age-group. Broadly, it was seen that the brain E2 slowly increased with age, peaking at 14 mos (**Fig. 4J**, p=0.062, F_3, 25_ = 2.77, one-way ANOVA). Brain P4 levels on the other hand fluctuated across the age-groups (**Fig. 4K**, p=0.001, F_3, 25 =_ 6.73, one-way ANOVA). P4 levels first decreased at 6 mos of age, then increased at 9 mos and then decreased again at 14 mos of age (**Fig. 4K)**. Spearman’s rank correlation was used to assess the relationship between hormone levels (E2 and P4) and the outcomes OD (at the 58:42 and 56:44 odor ratios) and DI in the OCPS task. There was a statistically significant positive association between log P4 and DI (p=0.010, **Fig. 4L**). Additional ANOVA results from linear regression, to determine effects of P4, E2 and age on DI for the OCPS task, confirmed that P4 and DI were significantly linked while E2 was not (**Fig. 4H**, p=0.04). Spearman correlation analysis showed that P4 was indeed significantly associated with DI (**Fig. 4I**, p=0.010).

### Ovariectomy causes drops in NSPC proliferation and neurogenesis at 9 mos of age

We utilized unbiased stereology by applying the optical fractionator probe via the MBF StereoInvestigator software to determine the number of NSPCs in the SVZ and DG labeled with the proliferation marker, BrdU, and DCX, a marker of neurogenesis. It was observed that the SVZ BrdU^+^ cells reduced with age, with a sharp decline at 14 months of age (**Fig. 5A-D and E**, p = 0.002 for 9 vs 14 mos, F_3,12_ = 15; one-way ANOVA with Tukey’s post-hoc test). On comparing Intact vs OVX groups at the different ages, a significant decline in SVZ BrdU^+^ cell number was seen at 9 mos of age (**Fig. 5F**, p = 0.009, F_1,24_ = 7.804 (group); two-way ANOVA with Tukey’s post-hoc test). A similar pattern of decline was seen with DCX^+^ neurons. DCX+ cells significantly decreased at the 14 mos aging stage (**Fig. 5G-J and K**, p = 0.001 for 9 vs 14 mos, F_3,12_ = 16.42; one-way ANOVA with Tukey’s post-hoc test), and in the 9 mos old OVX groups (**Fig. 5L**, p = 0.04, F_1,24_ = 10.62 (group); two-way ANOVA with Tukey’s post-hoc test). These data largely paralleled the SVZ relevant FOD behavioral data (**Fig. 1E-H**).

**Figure 5:**
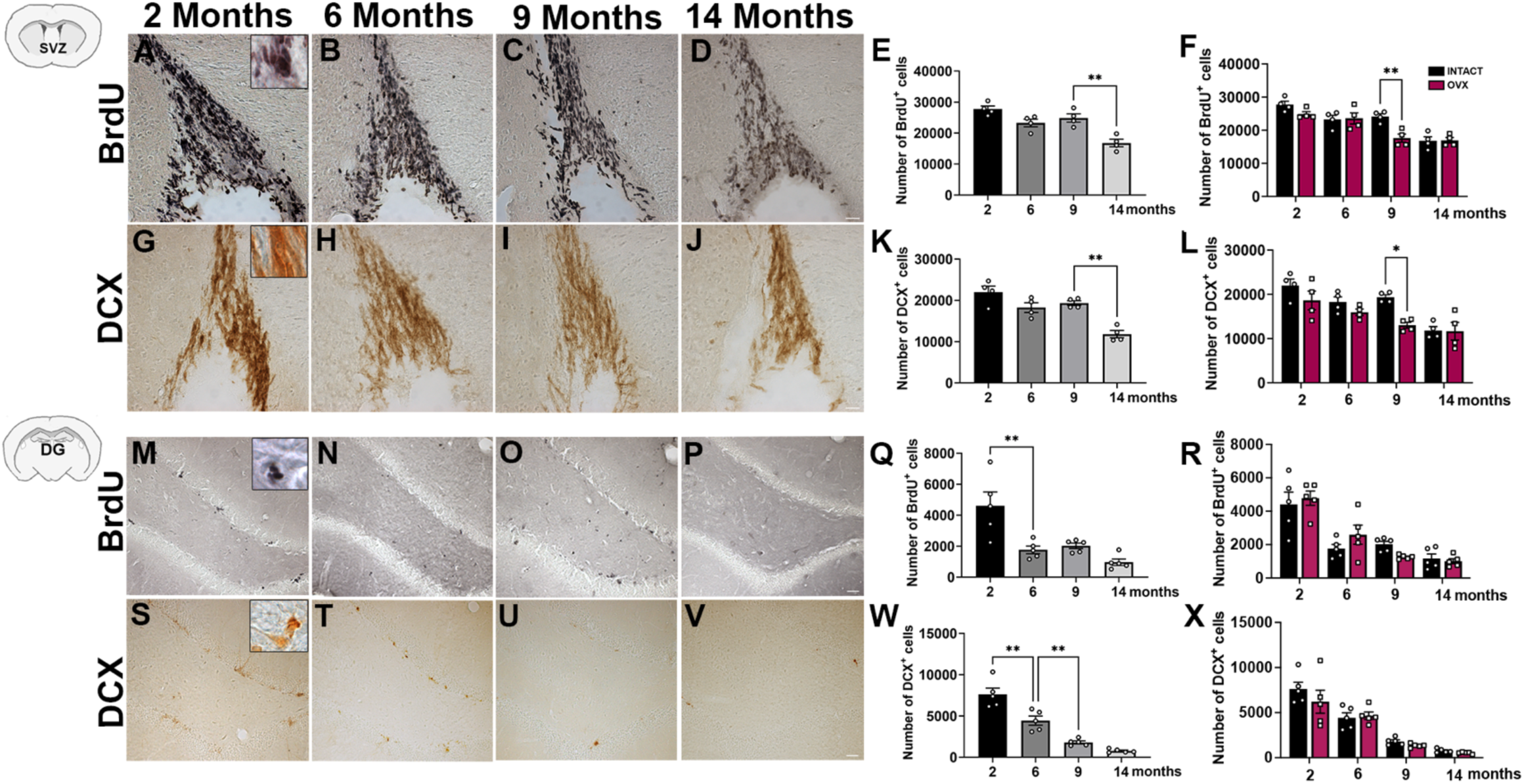
Comparison of proliferation and neurogenesis in the SVZ and DG during aging and after E2/P4 loss. (A-D) display representative images of BrdU immunostaining in the SVZ from 2, 6, 9 and 14 mos old Intact rats. Unbiased stereological quantification of the number of BrdU^+^ cells in Intact and OVX animals is in (E) and (F). (G-J) shows images of DCX immunostaining in the SVZ from the different age-groups with stereological quantification in (K) and (L) quantification. Similarly, results from the DG with respect to BrdU^+^ cells are in (M-R) and DCX+ cells is shown in (S-X). *** p = 0.001, ** p < 0.01, * p < 0.05; One-way ANOVA with Tukey’s post-hoc test (E, K, Q, W); Two-way ANOVA with Tukey’s post-hoc test and Unpaired t-tests (F, L, R, X); mean ± SEM, n = 4-5/group. Scale bar = 25μM.

With respect to DG NSPCs, the number of BrdU^+^ cells declined significantly already at 6 mos of age, and no other differences were seen subsequently (**Fig. 5M-P and Q**, p = 0.001 for 2 vs 6 mos, p = 0.006 for 6 vs 9 mos, F_3,16_ = 10.4; one-way ANOVA with Tukey’s post-hoc test). DCX^+^ cells similarly declined from at 6 mos of age, with an additional significant decline seen at 9 mos (**Fig. 5S-V and W**, p = 0.003 for 2 vs 6 mos, F_3,16_ = 41.7; one-way ANOVA with Tukey’s post-hoc test). When Intact vs OVX group differences at the different ages were analyzed, no changes in either BrdU or DCX^+^ cell numbers were seen (**Fig. 5R, X).** Interestingly, these data did not display similar trends as the OCPS and RMWM behavioral data, which showed significant differences between Intact and OVX groups at 9 and 14 mos of age. To explore this aspect further, we studied the morphology of DCX^+^ cells in the 4 age-groups (**Supp. Fig. 2**). Intact animals, especially at 9 and 14 mos of age, showed stronger DCX staining, and larger cells with evident neuritic extensions compared to DCX cells in the OVX group, which were smaller, mildly stained and had lesser, shorter processes (termed ‘distorted’) (**Fig. 2A-H**). Upon quantification, we found that these distorted cells tended to increase with age (p>0.05, One-way ANOVA, **Supp. Fig. 2I**). On comparing Intact and OVX rats, greater percentages of distorted DCX^+^ cells were seen in the 9 mos OVX rats (**Supp. Fig. 2J**, p=0.019, t=2.90, df = 8, Unpaired t-test). The 14 mos old OVX rats showed a tendency to harbor higher percentages of distorted DCX^+^ cells, but this was not statistically significant.

### The volume of the dorsolateral SVZ decreased with age and ovariectomy

The Cavalieri probe in the MBF StereoInvestigator software was used to assess whether age and/or OVX affected the volume of dorsolateral (DL) SVZ and the granule cell layer (GCL) of the DG (**Fig. 6A, D**). A significant decrease in the volume of DL SVZ was noted at 14 mos of age (**Fig. 6B**, p=0.015 for 6 vs 14 mos, F_3,16_ = 4.355, one-way ANOVA with Tukey’s post-hoc test). In terms of differences between Intact and OVX at each of the four age-groups, a significant decrease in the DL SVZ volume was seen after OVX at 6 mos (p = 0.01, t=3.88, df=8, Unpaired t test) while there were no differences at any of the other ages (**Fig. 6C**). In contrast, the estimated volume of DG-GCL, across 2, 6, 9 & 14 mos old rats did not show changes (**Fig. 6E**). Additionally, there were no Intact and OVX differences were noted within age-groups (**Fig. 6F**). These data indicated that while the volume of the DL SVZ declined with age and upon OVX at 6 mos, the DG-GCL volume remained stable.

**Fig. 6:**
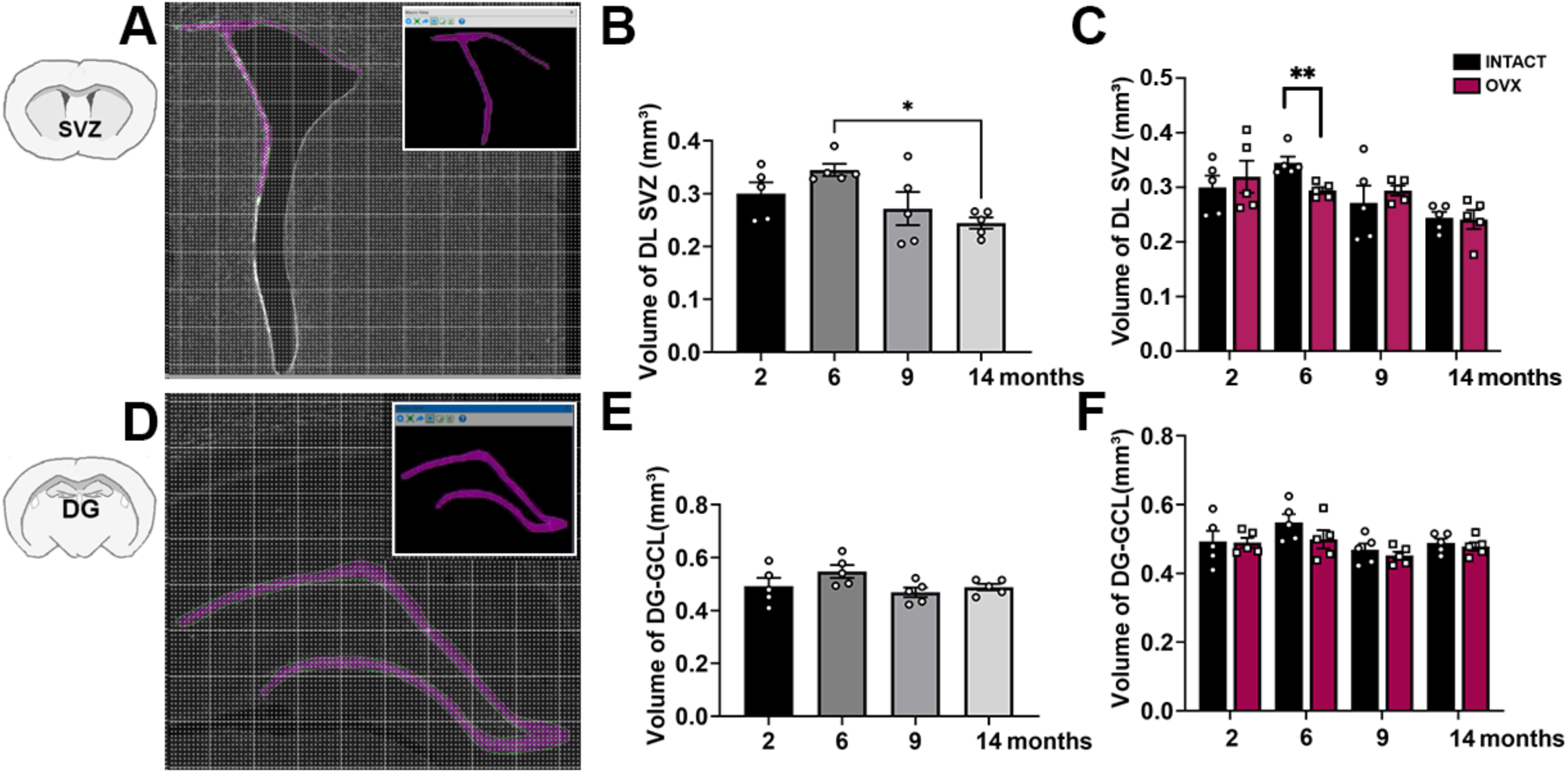
Volume changes in the SVZ and DG with age after ovariectomy induced E2/P4 loss. Cavalieri probe was applied to assess the volume of dorsolateral (DL) SVZ (A) and the granule cell layer (GCL) of DG (D) in the StereoInvestigator software. The insets in (A) and (D) show a magnified image of the contour with the Cavalieri probe markers in pink. (B) shows changes in the volume of SVZ across the 4 ages in Intact rats and the comparisons between Intact and OVX groups at each age is shown in (C). Similarly, volume estimations of the of the DG-GCL by age is in (E) and differences between Intact and OVX groups at each age is in (F). * p < 0.05; One-way ANOVA with Tukey’s post-hoc test (B, E); Two-way ANOVA with Tukey’s post-hoc test and Unpaired t-tests (C, F); mean ± SEM, n = 5/group.

### Ovariectomy affects the expression of estrogen and progesterone receptors in the SVZ and DG

We assessed the effects of age and ovariectomy on the expression of estrogen receptor alpha (ERα), estrogen receptor beta (ERβ) and progesterone receptor (PR) in the SVZ and DG using western blotting. A significant increase in ERα and ERβ expression was seen with age in the SVZ, particularly between 9 and 14 mos of age (**Fig. 7A, D**; ERα: p = 0.024, F_3,21_ = 8.04; ERβ: p = 0.0009, F_3,21_ = 13.5, one-way ANOVA with Tukey’s post-hoc test). Comparisons between Intact and OVX groups at the various ages in the SVZ showed a significant increase in ERα at 9 mos of age upon OVX (**Fig. 7B, example blots are in C**; p = 0.012, t=3.192, df=8, Unpaired t-tests). ERb also showed a similar increase at 9 mos after OVX (**Fig. 7E, F**; p = 0.025, t=2.744, df=8, Unpaired t-tests), although a decrease was also seen at 6 mos (**Fig. 7E, F**; p = 0.043, t=2.251, df=12, Unpaired t-tests).

**Figure 7:**
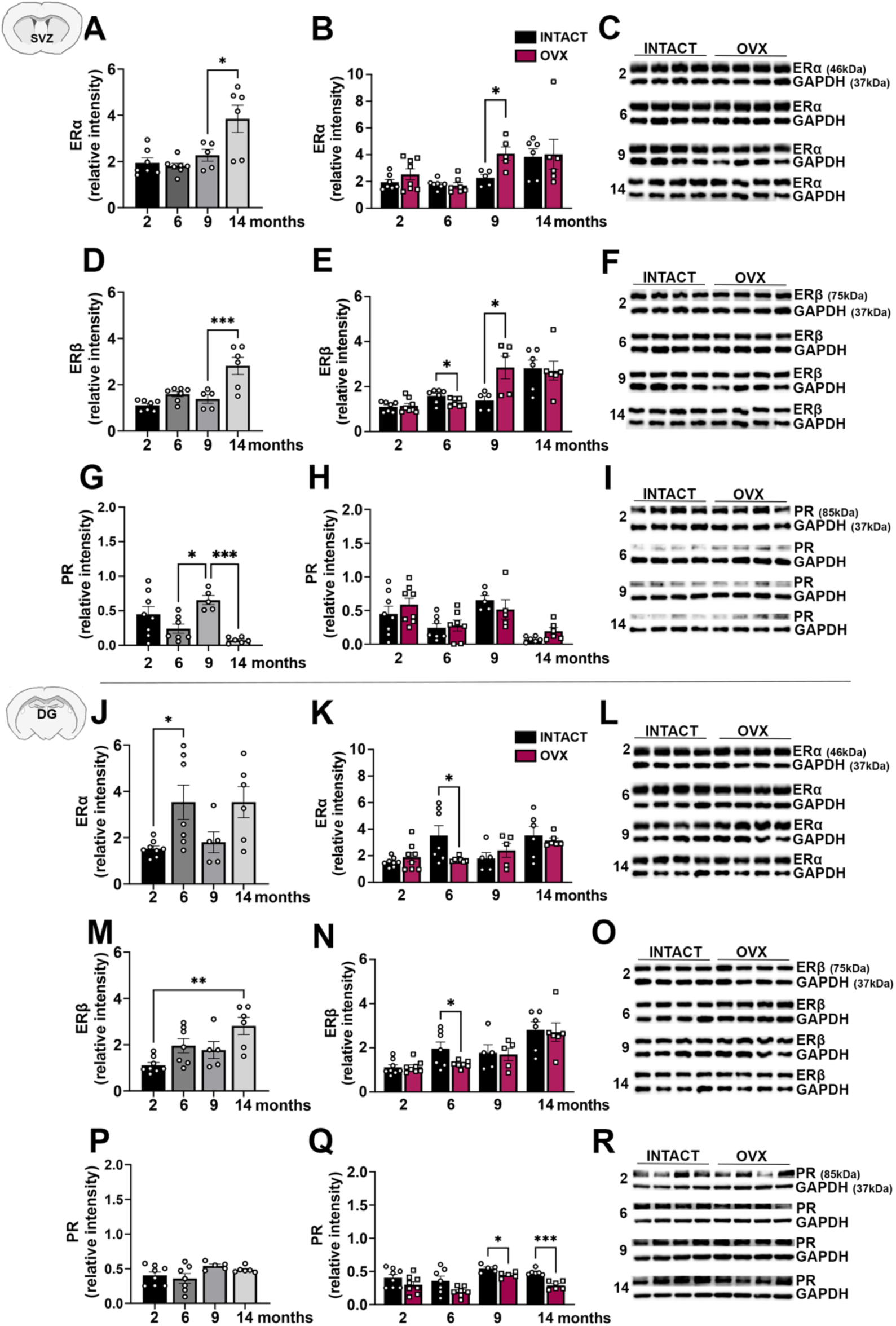
Alterations in estrogen and progesterone receptor expression in the SVZ and DG with age and acute E2/P4 loss. Quantification of ERα, ERβ and PR expression in SVZ tissues, across the 4 age-groups in Intact rats, as assessed via western blotting, are shown in (A, D, G). (B, E, H) show comparisons of ERα, ERβ and PR expression between Intact and OVX groups, with representative blot images in (C, F, I). Similarly, quantification of ERα, ERβ and PR expression in the DG, at the different ages is shown in (J, M, P). Comparisons of DG ERα, ERβ and PR expression between Intact and OVX groups is in (K, N, Q), with representative blot images in (L, O, R). *** p = 0.001, ** p < 0.01, * p < 0.05; One-way ANOVA with Tukey’s post-hoc test (A, D, G, J, M, P); Two-way ANOVA and Unpaired t-tests (B, E, H, K, N, Q); mean ± SEM, n = 6-8/group.

In the DG, a significant increase in ERα expression was seen between 2 and 6 mos of age, after which no statistically significant alterations were seen (**Fig. 7J**; p = 0.0249, F_7,44_ = 3.788, one-way ANOVA with Tukey’s post-hoc test). ERβ expression levels increased by 14 mos stage (**Fig. 7M**; p = 0.0005, F_7,44_ = 6.82, one-way ANOVA with Tukey’s post-hoc test). Both ERα and ERβ expression notably declined after OVX in the 6 mos old rats in comparison to Intact rats (**Fig. 7K, L**; ERα: p = 0.029, t=2.473, df=12; **Fig. 7N, O**; ERβ: p = 0.0411, t=2.28, df=12, Unpaired t-tests). No changes between Intact and OVX were seen at the other ages.

PR receptor expression increased significantly at 9 mos (p=0.01) but then reduced drastically at 14 mos of age (p=0.001) in the SVZ (F_3,22_ = 3.54, one-way ANOVA with Tukey’s post-hoc test), while its expression did not change with age in the DG (SVZ - **Fig. 7G, blot images in I; DG - Fig. 7P, blot images in R**). Intact versus OVX comparisons, at each of the four ages, showed no significant differences in the SVZ (**Fig. 7 H, blot images in I)**. In the DG, reduced PR receptor expression was found in the OVX group at 9 and 14 mos (**Fig. 7 Q, blot images in R;** 9 mos: p = 0.01, t=3.07, df=8; 14 mos: p = 0.001, t=6.20, df=10, Unpaired t-tests). These data indicated that ERα and ERβ expression generally increased with age and were altered upon OVX mainly at 9 mos in the SVZ and 6 mos in the DG. PR expression on the other hand, showed changes with age 9 and 14 mos in the SVZ (with age), and was reduced in the DG upon OVX at 9 and 14 mos of age.

### Age and ovariectomy alter antioxidant protein expression in the SVZ and DG

Increased stress due to a decline in antioxidant defenses and increased reactive oxygen species (ROS) is a known aspect of the aging process (Gemma et al., 2007; Ionescu-Tucker and Cotman, 2021; Schmidlin et al., 2019). Therefore, we examined the expression of NQO1, GCLM and HO-1, which are key ROS regulators/antioxidants. These molecules are also targets of the Nrf2 transcription factor, whose expression and activity are known to be altered during NSPC aging in males (Anandhan *et al*., 2021a; Corenblum *et al*., 2016; Dodson *et al*., 2021; Madhavan, 2015; Ray *et al*., 2018). In the SVZ, the expression of NQO1, GCLM, and HO-1 appeared to gradually increase with age (**Fig. 8A, D, G**). When the levels of NQO1 were compared between Intact and OVX rats no significant differences at any age were found (**Fig. 8B, sample blots in C**). GCLM on the other hand was downregulated in the OVX rats at 9 and 14 mos of age (**Fig. 8E, F**; 9 mos: p = 0.052, t=2.27, df=8; 14 mos: p = 0.004, t=3.64, df=10; Unpaired t-tests). HO-1 showed a similar trend to GCLM and was also reduced in the OVX animals at 9 and 14 mos (**Fig. 8H, F**; 9 mos: p = 0.022, t=2.81, df=8; 14 mos: p = 0.06, t=2.11, df=10; Unpaired t-tests).

**Figure 8:**
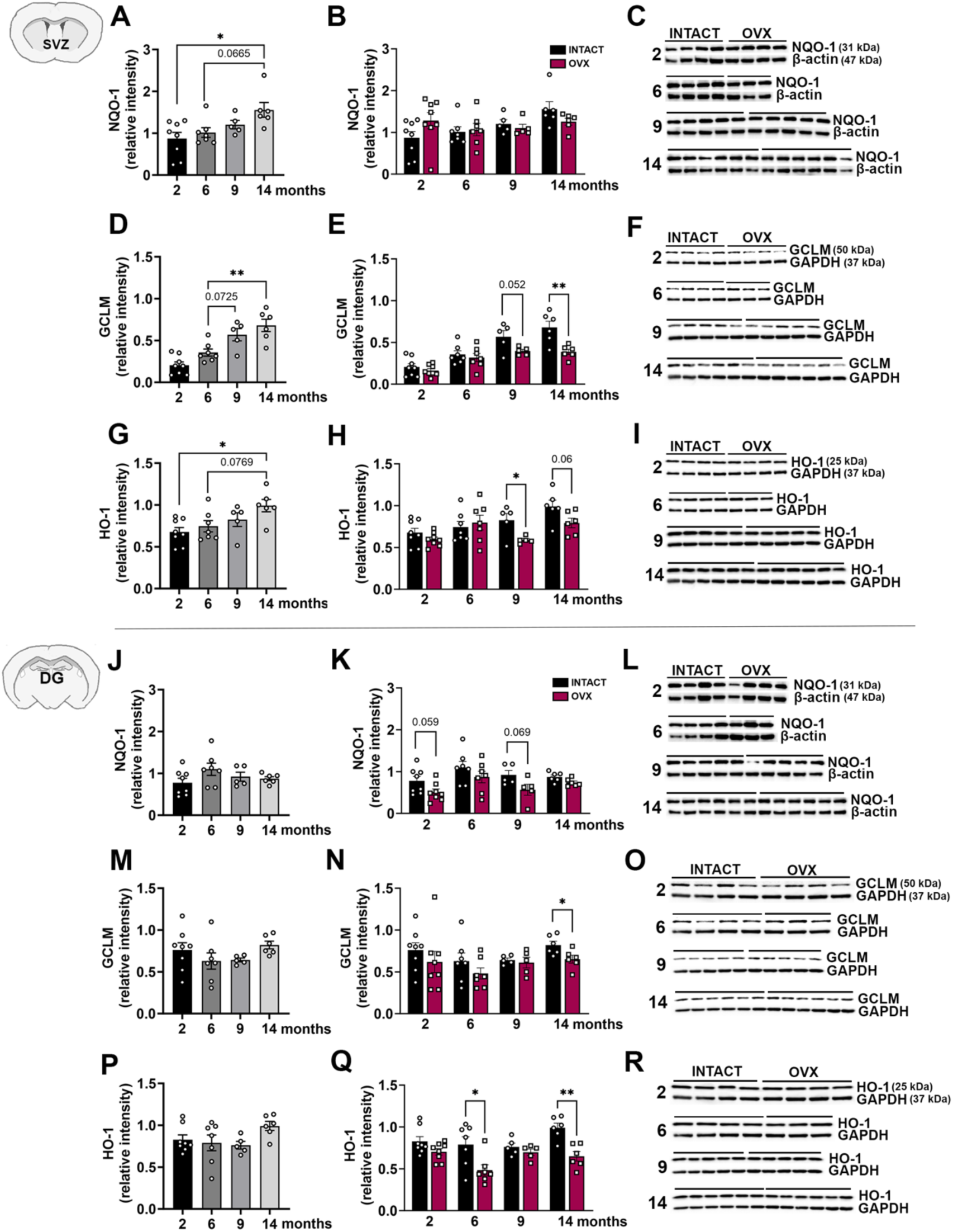
Age and ovariectomy affects antioxidant expression in the NSPC niches. Quantification of antioxidant protein expression (western blotting), specifically NQO-1, GCLM, and HO-1, in SVZ tissues in 2, 6, 9 and 14 mos old Intact female rats are shown in (A, D, G). (B, E, H) has comparisons of NQO-1, GCLM, and HO-1 expression between Intact and OVX groups, with representative blot images in (C, F, I). Similarly, quantification of NQO-1, GCLM, and HO-1 expression in the DG, with age (J, M, P). Comparisons of NQO-1, GCLM, and HO-1 expression in the DG between Intact and OVX groups appears in (K, N, Q), with representative blot images in (L, O, R). ** p < 0.01, * p < 0.05; One-way ANOVA with Tukey’s post-hoc test (A, D, G, J, M, P), Unpaired t-tests (B, E, H, K, N, Q); mean ± SEM, n = 6-8/group.

In the DG, no significant changes were noted in the levels NQO1, GCLM or HO-1 with age in the Intact rats (**Fig. 8J, M, P**). NQO1 expression was reduced at 2 mos and 9 mos of age in the OVX rats, with strong trends but no statistically significant effects (**Fig. 8K, sample blots in L**). OVX caused significantly lower GCLM expression at 14 mos (**Fig. 8N, sample blots in O;** 14 mos: p = 0.027, t=2.572, df=10; Unpaired t-tests), and lower HO-1 at 6 and 14 mos (**Fig. 8Q, sample blots in R;** 6 mos: p = 0.019, t=2.68, df=12; 14 mos: p = 0.001, t=4.35, df=10; Unpaired t-tests).

It is known that the elevation of AOX1, a detoxifying enzyme involved in the metabolism of various compounds, can augment ROS levels (Kucukgoze et al., 2017; Qiao et al., 2020). Thus, we also assessed AOX1 levels in the SVZ and DG. It was seen that with aging, AOX1 expression significantly reduced at 14 mos of age in the SVZ and increased at 9 mos of age in the DG (**Supp. Fig. 3A, D**; SVZ: 6 vs 14 mos, p = 0.048, F_3,44_ = 3.305; DG: 6 vs 14 mos, p = 0.024, F_3,22_ = 3.36, one-way ANOVA with Tukey’s post-hoc test). No changes were seen between Intact and OVX at the different ages in the SVZ (**Supp. Fig. 3B, sample blots in C).** Nevertheless, notable increases in the OVX rats were seen at 6 and 9 mos of age in the DG (**Supp. Fig. 3E, sample blots in F;** 6 mos: p = 0.015, t=3.07, df=8; 9 mos: p = 0.053, t=2.13, df=12; Unpaired t-tests.

## DISCUSSION

While the effects of age on neurogenesis have been studied in males, research in females is less abundant. From this perspective, our study captures the pattern of decline in neurogenesis in aging female F344 rats, pinpointing a vulnerable time at at 9 mos of age, when the acute loss of ovarian E2 and P4 further exaggerates this phenomenon. Behavioral deficits in olfactory discrimination, pattern separation and spatial learning were seen during this delicate time, underlining the functional relevance of the observed decline in neurogenesis. These deficits were accompanied by adaptive changes in the expression of estrogen and progesterone receptors, and antioxidants expression. To our knowledge, these results are the first to delineate how the stress of advancing age and sex hormone loss may affect the biology of adult neurogenic niches (in the SVZ and DG) of female rodents.

This study makes the following novel contributions. Firstly, it identifies a specific aging period when the acute loss of E2 and P4 makes the most impact on neurogenesis. Our data indicate that, at 9 mos of age, ovariectomy (OVX) compromises fine olfactory discrimination (FOD), a correlate of SVZ neurogenesis, and object-context pattern separation (OCPS) performance, a correlate of DG-based neurogenesis. These behavioral deficits were associated with OVX-mediated reductions in neurogenesis and/or distorted newborn neuron (DCX) morphology, thus defining a particular aging time that is sensitive to the loss of circulating E2/P4. Statistical analyses showed that the strongest effects of OVX in the 9 mos old animals occurred at the 58:42 and 60:40 odor ratios in the FOD task. Moreover, both the DI and % exploration of the novel object in the OCPS task were compromised in the 9 mos rats. Interestingly, a reduction in pattern separation ability was also seen in the 14 mos old OVX rats, although no statistically significant deficits in neurogenesis or DCX cell morphology were found in this group. This could suggest that the behavioral impairment at 14 mos was independent of neurogenesis.

Previous studies have investigated SVZ neurogenesis and olfactory discrimination in adult females in relation to learning male odors and pregnancy (Brus et al., 2016; Chaker et al., 2023; Puri et al., 2023). In terms of hippocampal neurogenesis, studies have reported that estradiol can improve pattern separation abilities in adult animals (Yagi et al., 2023). But less is known in the aging context. Our study systematically investigates age-relevant alterations in FOD, OCPS, and RMWM task performance in the presence or absence of circulating E2 and P4. Overall, the results indicate that the ovarian sex hormones may be particularly important to maintain neurogenesis between 6-9 mos of age. Importantly, the drops in neurogenesis also correlated to behavioral deficits. In the SVZ, a significant decline in FOD and neurogenesis was noted between 9-14 mos of age, while a more gradual decline in neurogenesis and spatial abilities was seen in the DG context. The RMWM task performance closely mirrored the changes in DG neurogenesis with significant decays at 6 mos of age after which progressive reductions happened till 14 mos. It is known that animals with reduced adult hippocampal neurogenesis, including in our own studies, show impairments in reversal learning (Garthe *et al*., 2014; Garthe and Kempermann, 2013). Specifically, DG neurogenesis can contribute to cognitive flexibility or the ability to adopt new strategies to complete a previously learned task when contingencies change – an ability that can be gauged through finding the platform location in the RMWM task (Anacker and Hen, 2017; Berdugo-Vega et al., 2021). Intriguingly, and contrary to our expectations, it was observed that the 9 mos old females that had undergone OVX were better at finding the location of the new platform on the first day of reversal (day 5), compared to the Intact rats. However, these 9 mos old OVX females also showed deficits in the acquisition phase of the task (first 4 days). Probe trial analysis conducted soon after the acquisition phase confirmed these findings and showed that the 9 mos old OVX animals in fact made significantly less goal crossings (Fig. 2B) and failed to form correct allocentric maps to find the correct location of the platform (Fig. 3F). This may have resulted in their ‘seemingly better’ performance on day 5 (reversal) to find the new platform location. In support, results showed that the 9 mos old OVX group used significantly more non-goal oriented and procedural (non-allocentric) search strategies, than allocentric strategies on day 4 (last day of the acquisition phase, Fig 2Q). In comparison, the 14 mos old Intact rats were better than their OVX counterparts in finding the new platform location on day 5 (Fig 2F) and showed higher allocentric confidence levels (Fig. 3F). On one hand, these data could mean that a certain availability of brain E2 and P4 was needed for NSPC functioning and proper reversal learning. On the other hand, these differences could be independent of neurogenesis especially given the lack differences in newborn neuron cell numbers or their morphology between the 14 mos old Intact and OVX rats. Future studies will assess these interesting findings.

Secondly, our study reports how estrogen and progesterone receptor expression change in the SVZ and DG niches, with aging and ovariectomy. With respect to the SVZ, a general increase in ERα and ERβ expression was seen with age, with a significant difference noted between 9-14 mos. Interestingly, this positively correlated with brain estrogen levels which increased at 14 mos of age. OVX amplified both ERα and ERβ expression at 9 mos of age, increasing them to 14 mos levels, signifying a compensatory response to the sudden loss of E2. In the DG, the ERα and ERβ receptor expression also increased with age although the pattern was different from the SVZ: ERα increased earlier at 6 mos of age while ERβ increased more gradually till 14 mos. Also, OVX altered receptor expression at 6 mos of age. Overall, post-OVX reductions were seen in the DG, compared to more elevations in the SVZ. These results may imply that receptor adaptations in response to circulating E2 loss maybe less efficient in the DG. Progesterone (PR) receptor expression differences were also seen between the SVZ and DG. For example, PR receptor expression was seen to fluctuate with age in the SVZ (decreases seen at 6 and 14 mos, and an increase at 9 mos). This pattern paralleled fluctuations in brain P4 levels. The DG was different in that no such PR receptor fluctuations were seen with advancing age.

We also investigated antioxidant expression with age and OVX, as a reflection of stress resistance in the SVZ and DG niches. Here, the data showed that antioxidant expression gradually increased across the age-groups in the SVZ. This change suggested an attempt to counteract increasing oxidative stress that is known to occur during aging (Finkel and Holbrook, 2000; Gemma *et al*., 2007). It was also seen that OVX notably reduced GCLM and HO-1 expression at 9 mos of age, correlating with the reductions in neurogenesis occurring in these animals. Reductions in GCLM and HO-1 were also seen in the 14 mos OVX group. As mentioned, these changes could be unconnected to neurogenesis which did not decline in upon OVX at 14 mos. In the DG, significant reductions in GCLM and HO-1 were mainly seen at 14 mos of age. These reductions could be contributing to the distorted newborn neuron morphology and behavioral compromises seen in OCPS and RMWM at this aging stage. These data are in line with studies that show that estrogen and progesterone are neuroprotective and modulate oxidative stress in the brain (Irwin et al., 2008; Nilsen, 2008; Razmara et al., 2007). These hormones are known to activate antioxidant defense systems scavenging ROS and limiting mitochondrial damage both in the context of normal aging as well pathology (Arnold and Beyer, 2009; Irwin *et al*., 2008; Nilsen, 2008; Razmara *et al*., 2007).

Finally, we report E2 and P4 concentrations in the brain tissues of aging females and characterize estrous cycle changes. Brain levels of E2 and P4 are not directly associated with ovarian function since E2 and P4 can also be produced at local sites in the brain. It is known that across vertebrates, the enzyme that synthesizes estrogens is expressed in several brain regions (Vahaba and Remage-Healey, 2015). Specifically, the brain can produce estrogen, known as brain-derived E2 (BDE2), primarily in neurons and astrocytes, through a process involving the enzyme aromatase, even in the absence of circulating ovarian hormones. Progesterone can also be synthesized locally in the nervous system by glia and neurons from cholesterol (Baulieu and Robel, 1990; Guennoun, 2020; Schumacher et al., 2014). Previous work has showed that BDE2 plays a key role for the maintaining of hippocampal neurogenesis and cognitive function (Huang et al., 2023). However, not much is known about how locally produced brain E2 and P4 may relate to neurogenesis, an area where our study makes an advance.

We also examined estrous cycle changes with age. Previous studies have looked at the estrous cycle in relation to neurogenesis in adult animals but have not addressed it in the context of aging. Our data show that the 2 and 6 mos old rats appear to be regularly cycling (Sone et al., 2007; Westwood, 2008). At 9 months, some of the rats continue to have regular cycles while others enter an extended estrus phase where the estrus stage is prolonged for 2-4 days instead of 24 hours. Most of the 14 mos old rats appear to have entered this extended estrus phase in our study. This aligns with other F344 rat studies that show a similar pattern of estrous cycle alterations with age (Sone *et al*., 2007). Furthermore, ANOVA results from linear regression indicated that estrous stage was not a significant predictor of either olfactory discrimination or pattern separation abilities (Fig 4I, J). Nevertheless, bigger sample sizes maybe needed to make clearer interpretations.

In conclusion, our study identifies a distinct female aging period, between 6-9 mos, during which SVZ and DG NSPCs are sensitive to the loss of circulating E2/P4. At the same time, the study also highlights specific differences between SVZ and DG neurogenesis and their associated biology. These findings are important in understanding the sex-specific regulation of the SVZ and DG stem cells, and how particular aging periods of reduced NSPC plasticity in females may engender the development of age-related disease. Previous studies have focused on adult neurogenesis mainly in male animals. It has been assumed that the findings in males naturally extend to females. The gamut of age-related changes in NSPC niches has also not been examined carefully and species differences are not well considered. This work advocates for carefully interpreting neurogenesis in the context of age, sex, and species differences, keeping with nature’s evolutionary diversity.

## MATERIALS AND METHODS

### Animals

Adult female Fisher 344 rats were obtained from the National Institutes of Health (NIH-NIA) and Charles River Laboratories and aged to 2, 6, 9 and 14 months. The animals were housed in the University of Arizona Animal Care Facility under a reverse 12-hour light-dark cycle condition with food and water available ad libitum. Animals were treated according to the rules and regulations of the NIH and Institutional Guidelines on the Care and Use of Animals. The University of Arizona Institutional Animal Care and Use Committee (IACUC) approved all experimental procedures.

#### Experimental Design

As shown in Figure 1A, rat cohorts at the different age-groups were first ovariectomized or underwent sham surgery. Two weeks after ovariectomy, the rats were subjected to different behavioral tests, relevant to NSPCs, namely fine olfactory discrimination (FOD), object-context pattern separation (OCPS), and reversal in the Morris water maze (RMWM) (Burghardt et al., 2012; Enwere et al., 2004; Goncalves et al., 2016; Kouremenou et al., 2020; Sahay et al., 2011). Subsequently, the animals were injected with 5′-bromo-2′-deoxyuridine (BrdU), to label proliferating cells, and sacrificed for downstream histological and molecular analyses. Vaginal smears were collected for estrous cycle staging after each behavior and just before sacrifice. The number of animals in each experimental group was as follows: 2 months - Intact (n = 16), OVX (n=16); 6 months - Intact (n = 15), OVX (n=15); 9 months - Intact (n = 13), OVX (n=13);14 months - Intact (n = 14), OVX (n=14).

#### Ovariectomy (OVX)

Rats were anaesthetized through intraperitoneal (IP) injections of Ketamine/Xylazine (70mg/kg and 10mg/kg respectively) until an appropriate plane of sedation was achieved. Subcutaneous analgesics were administered per IACUC protocols. All surgeries were performed under standard sterile conditions, and on a 35°C warming pad to prevent hypothermia. OVX was completed via one dorsal skin incision 2-3cm in length. The fascia lateral to the incision was dissected bluntly until the erector spinae muscle was visible. A small nick was made in the muscle to expose the ovary and fallopian tube. A hemostat was applied, and ligature placed over the fallopian tube and associated vessels medial to the hemostat. The fallopian tube and ovary were resected between the ligature and hemostat. The procedure was repeated on the contralateral side and the incision was then closed. OVX was later confirmed via vaginal cytology and E2/P4 measurements. A sham procedure was performed on the Intact group.

#### BrdU administration and labeling

Intraperitoneal 5′-Bromo-2-deoxyuridine (BrdU, Sigma-Aldrich, 50mg/kg, i.p.) was given twice daily, 12 hours apart, for the 3 days prior to sacrifice to label NSPCs (Anandhan *et al*., 2021a; Corenblum *et al*., 2016; Madhavan et al., 2015; Madhavan et al., 2012).

#### Euthanasia and post-processing

Animals were sacrificed via pentobarbital (60 mg/kg) overdose and perfused with 0.9% Saline. For histology, the animals were perfused with 4% paraformaldehyde (PFA; Electron Microscopy Sciences), brains were extracted, post-fixed in 4% PFA solution, sunk through a 30% sucrose solution, and coronally sectioned at 35 microns on a freezing stage sliding microtome. Brains to be used for molecular analysis were immediately flash frozen after saline perfusion and stored at −80C.

### Behavioral tests

#### Fine olfactory discrimination (FOD)

Rats were subjected to a fine olfactory discrimination task (FOD), as we have previously described (Anandhan *et al*., 2021a; Corenblum *et al*., 2016). Briefly, after a 2-day water deprivation, the rats underwent training and subsequent testing for olfactory discrimination abilities. For the training stage, 12 μl of double distilled water was placed in a sterile dish, and 1 μl of coconut extract (COC) was added. This combination served as a reward for response and was designated as positive [+]. The dish was placed at one end of the cage, and the rat was allowed 2 min to find and drink the [+]. Once the rat finished drinking, the dish was immediately removed and replaced with a fresh [+] solution after a 30 sec inter-trial interval. The amount of COC was increased with each trial until it reached 8.5 μl per dish per trial. From here, five additional trials were conducted using [+]. For the sixth trial, we presented the rats with a dish containing 8.5 μl of almond extract (ALM) added to 12 μl of a 1% solution of denatonium benzoate (DB: Sigma-Aldrich). This combination of ALM and DB was designated as negative [−] as the rats learned to associate the bitter taste of DB with the smell of ALM and avoided drinking the [−]. Four additional trials were conducted with [−] to ensure that the rats had learned to avoid the [−]. During the fine discrimination testing phase, the pure COC [+] and ALM [−] were replaced with a nearer proportions of both COC and ALM to produce a graph of performance against odor concentration, known as a generalization gradient. Rats were required to discriminate between solutions containing [+] ratios of COC:ALM of 100:0 (discrete discrimination), 60:40, 58:42, and 56:44 (fine discrimination) as well as the corresponding [−] ratios of COC:ALM of 0:100, 40:60, 42:58 and 44:56. The first trial utilizing the [+] mixture and its corresponding [−] mixture was conducted with the same 2 min time limit and 30 sed inter-trial interval. This was repeated so that five total trials were conducted for each combination of odorant. The COC and ALM dish locations were randomized within the cage throughout-testing.

#### Object-Context Pattern Separation (OCPS)

As described in our previous work, rats were first habituated to the testing room (Ray *et al*., 2018). Then, during the training period, they were placed in an arena (30 cm length × 30 cm width x 30 cm high) with black walls and floor and allowed to explore for 5 min. After a 30 min interval, animals were placed in the arena now containing a specific floor pattern and two identical objects, and were allowed to explore for 10 min. Following a 30 min inter-trial interval, animals were placed back in the arena now containing a different floor pattern and two new identical objects unique from the objects in the first trial and were again allowed to explore for 10 min. After 3 hours of memory incorporation, the testing period was started during which the animals were placed in the arena for 10 min with the floor pattern from trial two, one object from trial one, and one object from trial two. Time spent exploring the novel object (i.e. the object from trial one in the context from trial two) was compared with the time spent exploring the familiar object (i.e. the object from trial two in the context from trial two). The exploration time of each object was scored by a blinded individual from captured videos and was defined as the length of time the animal spent actively interacting with the object (when the rat’s nose was 1 cm away from the object) or the rat’s paws were touching the object. The Discrimination Index (DI) was calculated as: (total time with novel object minus total time with familiar object)/(total time spent exploring both objects).

#### Reversal in the Morris water maze (RMWM)

As described by Ray et al., (Ray *et al*., 2018), animals were tested in a circular pool of water (183 cm in diameter) made opaque by the addition of non-toxic tempera powdered paint, containing a submerged platform and fixed visual cues around the room. All trials were recorded with a video camera placed above the center of the tank and ANY-maze software (Stoelting Co., Wood Dale, IL, USA) was used to run trials and calculate various outcome measures. Testing occurred over a period of 6 days, in which the first 4 days encompassed a spatial task where the rats were dropped in the pool at varying locations to learn the location of the hidden escape platform (acquisition phase). A total of six trials per day were completed, with a rest period between every block of two trials. At the end of day 4 of the water maze, a probe trial was conducted where the hidden platform is removed altogether. Focal searching behavior in the probe trial was assessed through quantification of the number of goal crossings (previous location of the platform) or time spent in the appropriate quadrant of the removed platform. On days 5 and 6, the platform was moved 180 degrees in the opposite direction from its original location, and another six trials per day divided into blocks of two trials were performed for the additional 2 days. This constituted the reversal portion of the task (reversal phase).

#### Strategy Analysis in RMWM

The specific search strategy used on each RMWM trial was estimated using the approach described by Garthe et al. (Garthe *et al*., 2009) and implemented in the RTrack software package (Overall et al., 2020). RTrack performs an automated algorithm-based classification of MWM behavior based on the xy-coordinates of the animal in each trial and computes the likelihood the swimming behavior falls into one of 9 categories: direct path, directed search, corrected path, thigmotaxis, circling, random path, direct path, directed search, and corrected path. While details of the algorithm are described in Garthe et al. (Garthe *et al*., 2009), the categorization is based on factors such as percent-time on a trajectory approaching the goal, percent of pool surface coverage, average heading error, etc. The 9 search strategies can be grouped more broadly into allocentric, procedural, and escape strategies as shown in Figure 2G and described in (Overall *et al*., 2020). For the analysis of strategy preference during MWM acquisition (days 1-4), a two-way repeated measures ANOVA was performed (within = day, between = OVX/Intact) where the outcome variable was the likelihood of a given strategy. This likelihood the animal was using each strategy on a given day of testing was determined by averaging likelihoods for each strategy over the 6 within-day trials. This produced a single daily estimate of the likelihood for each strategy for each animal. Analysis of reversal performance was restricted to Day 5 and analyzed by averaging the likelihood of the target strategy (e.g., allocentric) across the 6 trials to produce a single likelihood for each animal. Group differences were tested using a t-test. Data analysis code in R is available on Github at https://github.com/CowenLab/Ovariectomy_and_development.

### Vaginal Cytology

Vaginal smears were prepared from each Intact animal following each behavior task and just prior to sacrifice. Vaginal samples were also collected 10 days after ovariectomy, at a consistent time each day, for four consecutive days, to confirm that the animals had stopped cycling. Briefly, a solvent-resistant permanent marker was used to draw a grid on glass microscope slides so that all smears for a particular animal were on one slide for each set of lavages. Using a micropipette, approximately 100µL of sterile water was flushed into the vagina 2-3 times ensuring the end of the pipet did not go into the canal itself. The fluid was dropped onto the prepared gridded slide and allowed to dry overnight at room temperature (RT). After each grid was completed and dried, the slides were stained with 0.1% Crystal Violet solution for 1 min, then washed with distilled water twice for 1 min each to remove excess stain. The slides were allowed to dry at room temperature for 1-2 hours after which they were ready for staging.

The reproductive phase of the rats was determined by the cell cytology in the vaginal smear (Fig 4A). Using light microscopy, the relative ratio of cell types (large and small nucleated epithelial cells, cornified epithelial cells and leukocytes) in the vaginal smear was assessed to classify the estrous stage the animal was in at the time of smear collection. The stage dominated with clusters of nucleated epithelial cells was identified as Proestrus. Estrus stage was recognized by smears with primarily cornified epithelial cells. A stage chiefly of leukocytes with some cornified epithelial cells was identified as Metestrus, and smears dominated by leukocytes with some nucleated epithelial cells or cluster formation was classified as Diestrus.

### E2 and P4 levels

LC-MS/MS was applied to measure estradiol and progesterone in brain tissues (Wisconsin National Primate Research Center) (Kenealy et al., 2013; Skalski et al., 2024). To extract and measure progesterone and estradiol, samples were homogenized in methanol containing internal standard using a sterile disposable pestle (DWK Life Sciences, Rockwood, TN). The samples were further homogenized using a MM400 ball mill (Retsch, Haan, Germany) operating at 30hz in three sessions of 5 minutes for a total of 15 minutes. They were homogenized a third time using a Polytron PT 10-35 homogenizer. The homogenate was cooled at −20°C for 30 minutes then centrifuged at 12,000 g for 5 minutes at 4°C. The resulting supernatant was evaporated to dryness and resuspended in 5% methanol for further purification. Steroid extraction and sample cleanup was achieved using an Oasis HLB solid phase extraction (Waters Corporation). After cartridge conditioning and equilibration, the samples were loaded onto the cartridge and washed with 20% methanol. The steroids were eluted from the cartridge using 90% methanol. The final eluate was evaporated to dryness and derivatized with dansyl-chloride in acetone. The samples were then analyzed by liquid chromatography-tandem mass spectrometry (LC-MS/MS) using a QTRAP 6500+ triple quadrupole (Sciex) adapted from previously published methods (Kenealy et al., 2013; Skalski et al., 2024). Intra and inter-assay CV was assessed with a pool of mouse hippocampi and ranged from 1.7 to 8.7 and 8.9 to 19.3%, respectively.

### Immunohistochemistry

Immunohistochemistry was performed according to standard protocols as described previously (Anandhan *et al*., 2021a; Anandhan et al., 2021b; Corenblum et al., 2015; Corenblum *et al*., 2016). Free floating tissue sections were washed in 1X Tris-Buffered Saline (TBS, pH 7.4), subjected to antigen retrieval in boiling Citrate buffer (1.8 mM citric acid, 8.2 mM sodium citrate, pH 6.0) if needed, and subsequently treated with blocking solution (10% normal goat serum, 0.5% Triton-X-100 in TBS) for 2 hours. With respect to BrdU immunostaining, sections were first treated with 2N HCL for 15 minutes at 37°C followed by 3 x 5 minutes washes in 0.1M Borate Buffer. Sections were then incubated overnight in the appropriate concentration of primary antibodies (see antibody table in Supplementary section for more details). For chromogenic detection, sections were pretreated with 0.3% Hydrogen peroxide for 30 min at 37°C, washed in 1X TBS, blocked for 2 hours and incubated overnight in primary antibodies. After washing in TBS, sections were incubated with biotinylated secondary antibodies for 2 hours, washed again, exposed to Avidin-Biotin-Complex (Elite ABC Kit, Vector Laboratories) for 2 hours, and signal was detected with 3,3′-Diaminobenzidine (DAB, Sigma-Aldrich) or a combination of DAB and Ammonium Nickel Sulfate (Ni, Sigma-Aldrich). All incubations were performed at RT unless noted otherwise. The following primary antibodies were used: Doublecortin (DCX; 1:1000, Abcam); BrdU (1:100, Abcam).

### Immunoblotting

Proteins were extracted from flash frozen brain tissue using RIPA buffer (50 mM Tris–HCl, 150 mM NaCl, 0.1% SDS, 0.5% sodium deoxycholate, 1% Triton X-100, pH 7.4) supplemented with protease inhibitor cocktail (1:100, MilliporeSigma, Burlington MA), phosphatase inhibitor Sodium Fluoride (1mM, Thermo Fisher Scientific), and protein tyrosine phosphatase inhibitor Sodium Orthovanadate (1mM, MilliporeSigma). Protein concentration was determined using a standard bicinchoninic acid (BCA) assay. A total of 10–50 µg of protein was separated by electrophoresis on a Mini-Protean Tetra Cell system using 7.5% and 10% TGX polyacrylamide gels (Bio-Rad, Hercules CA), followed by transfer (Trans-Blot Turbo, Bio-Rad) to 0.22 and 0.45µm PVDF membranes (MilliporeSigma). Membranes were blocked in 5% non-fat dry milk (Bio-Rad) in TBST (Tris-buffered saline with 0.1% Tween 20) or 5% Bovine Serum Albumin (BSA) (Fischer BioReagents) in TBST and incubated overnight with primary antibodies diluted in Hikari 250 Primary Antibody Signal Enhancer (Nacalai USA, San Diego, CA) at 4°C with gentle agitation. Subsequently, membranes were incubated with the appropriate horseradish peroxidase-conjugated secondary antibody for 1-2 hours at RT and visualized using Super Signal Femto maximum sensitivity chemiluminescent substrate (Thermo Fisher) on an Azure 500 imaging system (Azure Biosystems, Dublin CA). Optical density was quantified using Image Studio Lite software (LI-COR Biosciences, Lincoln NE), and protein levels were normalized to GAPDH or β-actin. The following antibodies were used: AOX1 (1:500; Proteintech), NQO-1 (1:1000; Santa Cruz Biotechnology), HO-1 (1:500; Santa Cruz Biotechnology), GCLM (1:500; Santa Cruz Biotechnology), Estrogen Receptor alpha (1:3000; R&D), Estrogen Receptor beta (1:5000; Invitrogen), Progesterone Receptor (1:500; Invitrogen), GAPDH (1:2000; Cell Signaling), β-actin (1:1000; Santa Cruz Biotechnology). More information can be found in the Supplementary section.

### Stereology

#### BrdU and DCX cell counts

Stereological probes were applied using a Zeiss Imager M2 microscope (Carl Zeiss, Oberkochen Germany) equipped with StereoInvestigator software (v2019.1.3; MBF Bioscience, Williston VT), according to our published methods (Anandhan *et al*., 2021a; Corenblum *et al*., 2016; Madhavan *et al*., 2012). Using the optical fractionator workflow, BrdU and Doublecortin (DCX) cell counts were conducted through the dorsolateral SVZ in sections at 480 μm intervals across the rostrocaudal axis of the structure. In all cases, after section thickness was determined, guard zones were set at 2 μm each at the top and bottom of the section. All contours were drawn around the region of interest at 10X magnification. Clear uniformly labeled BrdU nuclei or DCX positive cells were counted under a 63X oil immersion objective. The grid size and counting frame for BrdU and DCX counts in the SVZ were 65 x 65 μm and 50 x 50 μm respectively, and for the DG were 100 × 100 μm; counting frame was 80 x 80 μm, respectively. The counting frame was lowered at 1–2 μm interludes and each cell in focus was marked. Coefficients of error (CE) were calculated via the Gundersen method (m=1) and used to estimate the accuracy of the optical fractionator results. Coefficients obtained were generally less than 0.10.

#### Volume estimation via Cavalieri

The Cavalieri principle was used to estimate the volume of the dorsolateral SVZ (DL SVZ) and granule cell layer (GCL) of the DG. Briefly, the Cavalieri probe was applied on DAPI counterstained sections for unbiased estimation of volume through the StereoInvestigator software. For the ROI volume estimation, variable number of sections per animals were used from the most rostral section where the region was identified till the appearance of 3^rd^ ventricle for DL SVZ and the lateralization of DG for DG-GCL. Every 12^th^ section with a section thickness of 35 μm was assessed, ROI’s traced, and the Cavalieri probe applied to fill markers into the contour with a grid spacing of 25 μm. The volume was calculated by the formula Volume of a structure = (Total number of points marked * Distance between points in XY) * (Distance between points in Z). Volume = Σ (Area * Thickness).

### Statistical analysis

GraphPad Prism 10 software (San Diego, CA, USA) was used for statistical analyses. For comparisons between three or more groups, one-way analysis of variance (ANOVA) followed by Tukey’s or Bonferroni’s post-hoc test for multiple comparisons between treatment groups was conducted. Unpaired t-tests were used for comparisons between two groups. For analyzing the CIPL scores on the RMWM task, a two-way repeated measures ANOVA followed by a Šídák’s multiple comparisons test was used. Differences were considered significant at p < 0.05, and results are reported as mean ± SEM. Statistical details pertinent to each experiment are provided within the relevant results and legend sections.

For Figure 4, linear models were used to assess whether age, estrous stage or hormone levels were significantly associated with olfactory discrimination and pattern separation. Hormone levels were log-transformed for statistical analysis to improve symmetry of distributions. Results were interpreted by inspection of coefficients and associated hypothesis tests, as well as analysis of variance. Analyses were performed using R Studio version 2024.12.1.

## ACKNOWLEDGEMENTS

We thank Dr Akira Uruno (Tohuku University) and Dr Keiko Taguchi (Tokyo University) for their guidance with probing the antioxidant proteins via western blotting. The brain E2/P4 levels were measured at the Assay Services lab at the Wisconsin National Primate Research Center supported by P51OD011106 and S10OD028626, and we thank Amita Kapoor and Cody Corbett in this regard. We acknowledge resources from the UA Evelyn F McKnight Institute to perform the water maze experiments. The study was supported by a National Science Foundation grant (IOS 2207023) and UA Intramural funds to LM, Department of Physiology Teaching Assistantship to MS, NIH post-doctoral training grant (T32 AG044402) to AI, UA UBRP program support for TM, and UA FRONTERA and MARC program support for PW.

## AUTHOR CONTRIBUTIONS

SVP - Experimental design, Collection and assembly of data, Data analysis and Interpretation, Manuscript writing

MJS – Experimental design, Collection and assembly of data, Data analysis and Interpretation, Manuscript writing

MJC - Collection and assembly of data, Data analysis and Interpretation, General technical support, Manuscript editing

AI - Collection and assembly of data, Data analysis and Interpretation, Manuscript editing

TM – Collection and assembly of data, Data analysis

PW – Collection and assembly of data, Data analysis

NM – Collection and assembly of behavioral data, Data analysis

DB – Statistical correlation data analysis and interpretation, Manuscript editing

JK – Statistical correlation analysis

GW – Analysis of behavioral data

SC – Conceptualization and analysis of behavioral data

LM – Overall conception and design of the study, Data analysis and Interpretation, Manuscript writing, Financial support, Final approval of manuscript.

## SUPPLEMENTARY SECTION

### Western Blotting

A single membrane was used to detect various combinations of target proteins, including HO-1 and GCLM. These, along with GAPDH used as a loading control, were obtained from the same membrane via re-probing. AOX1 and NQO-1 were probed on separate membranes with β-actin as a loading control. Estrogen Receptor alpha and Estrogen Receptor beta were probed for sequentially on the same membrane without stripping with GAPDH as the reference standard. Progesterone Receptor was probed by stripping of membranes previously probed for GCLM and HO-1. Full blots can be displayed as requested.

**Supplemental Table 1:** Listing of antibody details in relation to the western blot and immunohistochemical studies.

| ANTIBODY | VENDOR | CATALOG # | HOST/TYPE/CLONE | CONCENTRATION |
| --- | --- | --- | --- | --- |
| Anti-Heme Oxygenase 1 (HO-1) | Santa Cruz Technologies | sc-390991 | Mouse/mAb/F-4 | 1:500 WB |
| Anti-Gamma-GCSm (GCLM) | Santa Cruz Technologies | sc-55586 | Mouse/mAb/E-4 | 1:500 WB |
| Anti-NQO1 | Abcam | ab2346 | Goat/pAb/C-terminus | 1:1000 WB |
| Anti-Aldehyde Oxidase 1 (AOX1) | Proteintech | 19495-1-AP | Rabbit/pAb | 1:500 WB |
| Anti-GAPDH | Cell Signaling Technology | 5174 | Rabbit/mAb/D16H11 | 1:2000 WB |
| Anti-Beta-Actin | Santa Cruz Technologies | sc-47778 | Mouse/mAb/C4 | 1:1000 WB |
| Anti-Estrogen Receptor Alpha (ERa) | R&D Systems | MAB57151 | Mouse/mAb/NR3A1 | 1:3000 WB |
| Anti-Estrogen Receptor Beta (ERb) | Invitrogen | PA1-311 | Rabbit/pAb/NR3A2 | 1:5000 WB |
| Anti-Progesterone Receptor | Invitrogen | MA1410 | Mouse/mAb/MA1-410 | 1:500 WB |
| Anti-Bromodeoxyuridine (BrdU) | Abcam | ab6326 | Rat/mAb/BU1/75 | 1:100 IHC |
| Anti-Doublecortin (DCX) | Cell Signaling Technology | 4604 | Rabbit/pAb | 1:500 IHC |

## SUPPLEMENTAL FIGURES

**Supplemental Figure 1:**
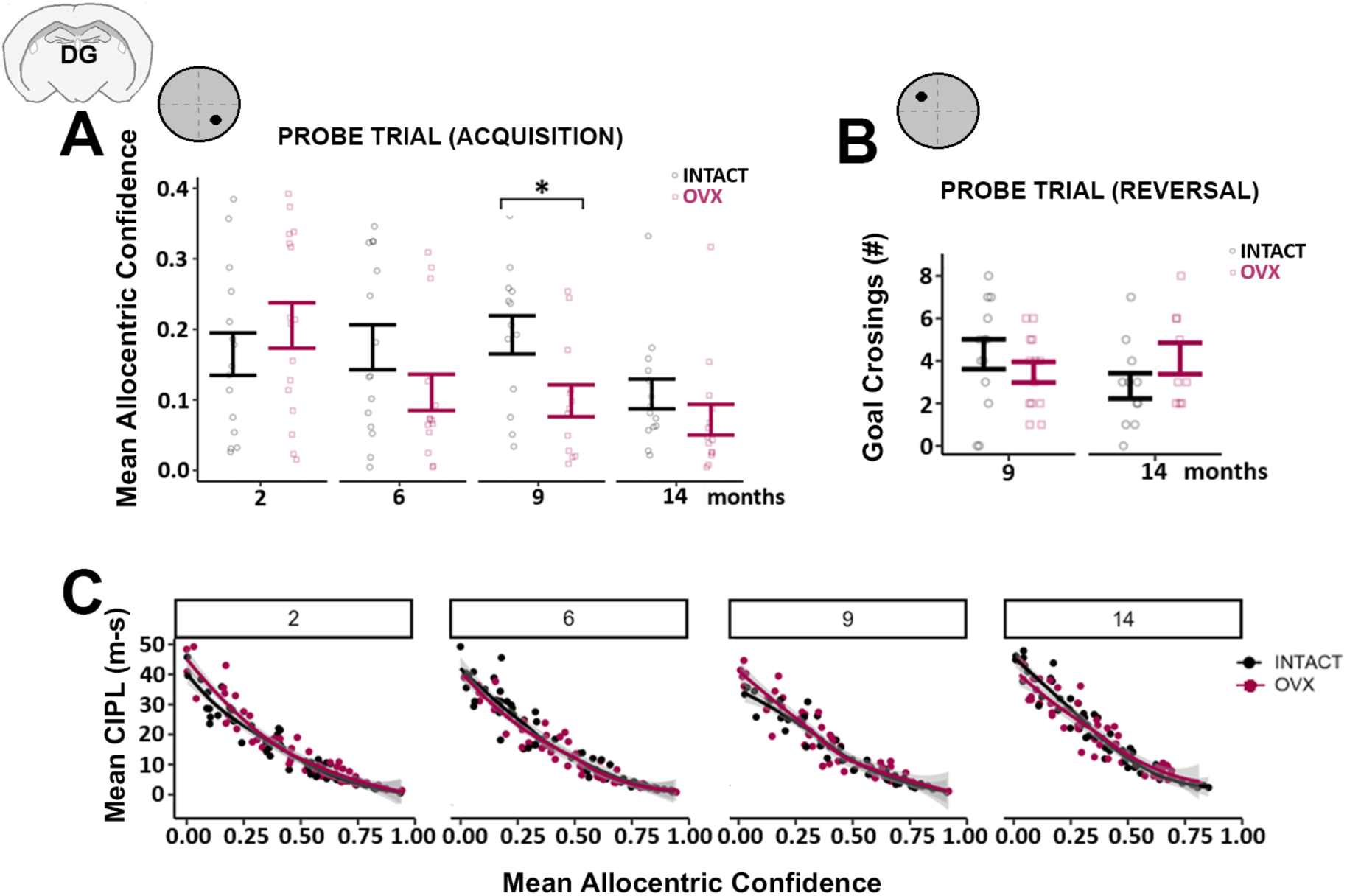
(A) The likelihood animals were using the allocentric strategy (Allocentric Confidence) during the probe trial identified reduced use of allocentric strategies in OVX animals relative to Intact in the 9 mos group (p < 0.05, t-test with Holm-Bonferroni correction). (B) Shows results (number of goal crossings) from the probe trial after the reversal phase, from the 9 and 14 mos old animals. No difference between groups was identified. (C) Scatterplots demonstrating the strong negative correlation between CIPL scores (y axis) and allocentric confidence (x axis) at each age group. A local regression line (Loess using R) was fit to the data. *p<0.05, two way RM-ANOVA, mean ± SEM, n = 13-16/group, t-tests were used for post-hoc comparisons using the Holm-Bonferroni correction for multiple comparisons.

**Supplemental Figure 2:**
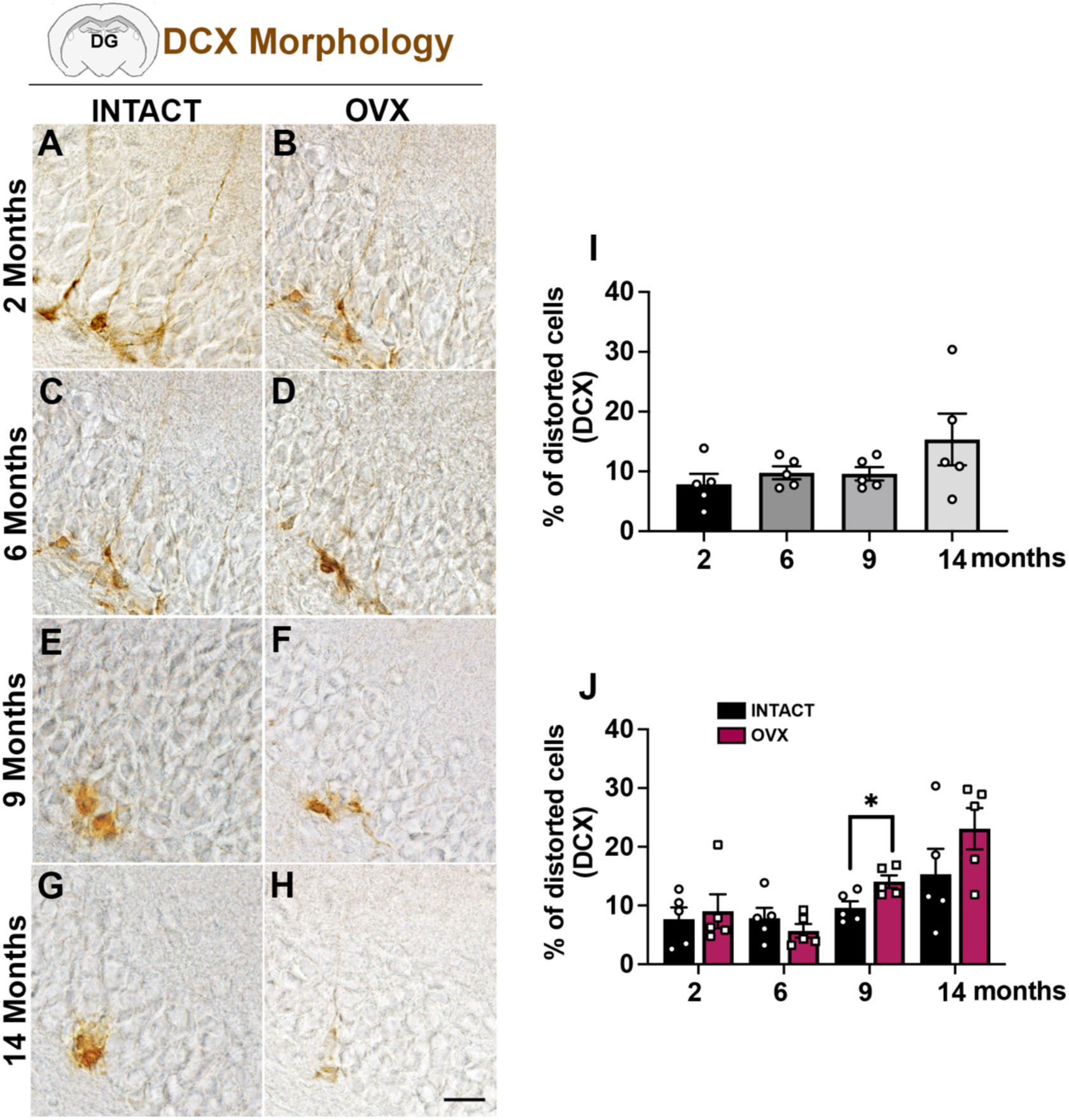
The morphology of newborn neuroblasts is compromised by age and ovariectomy. Representative DCX^+^ cells in the DG of Intact and OVX rats at 2 (A, B), 6 (D, E), 9 (G, H) and 14 (J, K) months of age are shown. Percentage of distorted DCX^+^ cells (mild staining, small soma and short processes) cells was quantified and shows that they generally increased with age (I). Comparison of distorted DCX^+^ cell percentages across Intact vs OVX groups (J). *p<0.05, One-way ANOVA with Tukey’s test in I; Unpaired t tests in J. Scale bar = 25μM.

**Supplemental Figure 3:**
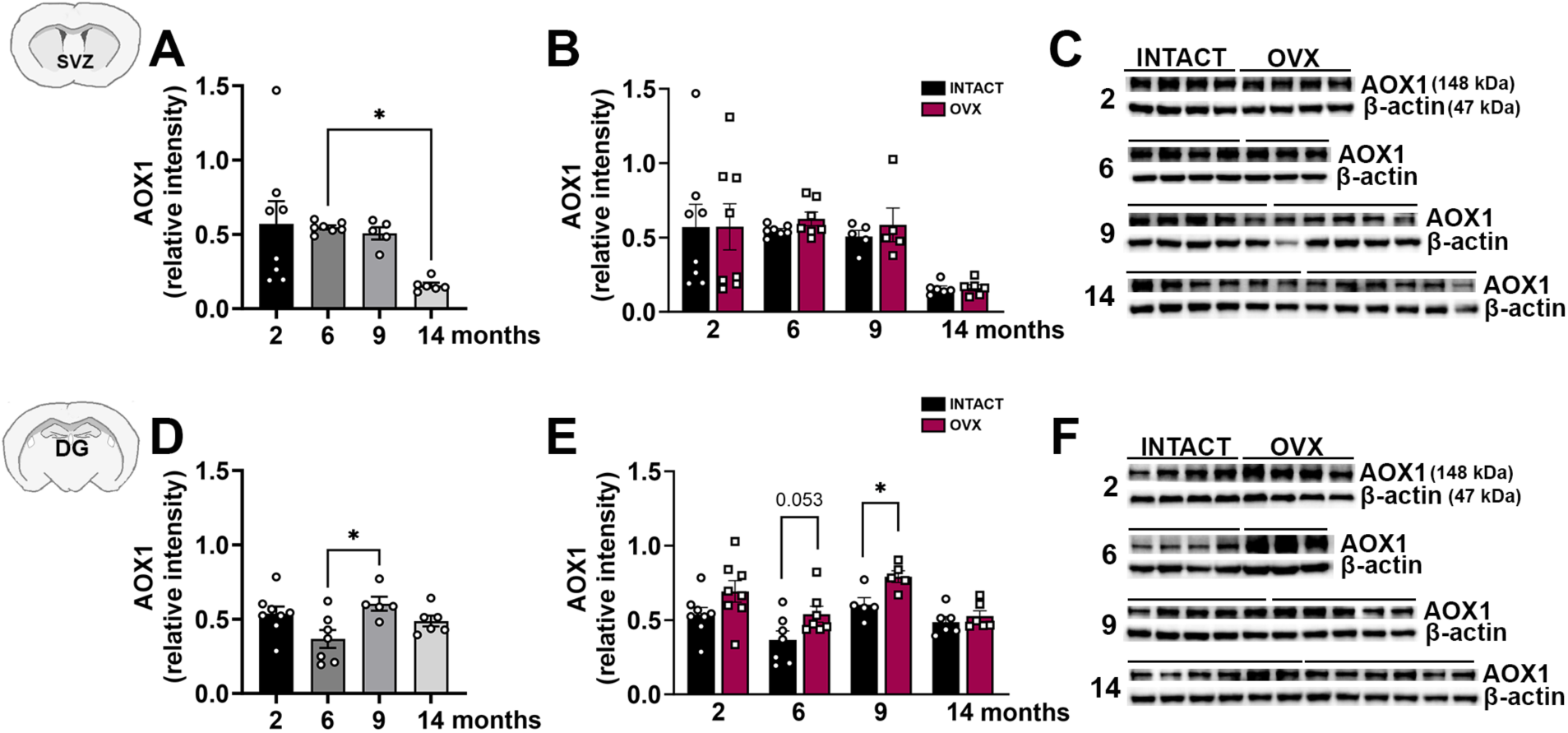
Effect of age and ovariectomy on AOX1. Quantification of the relative expression of the ROS modulator, AOX1 (western blotting), in the SVZ and DG of 2, 6, 9 and 14 mos old female rats is shown in (A, D). (B, E) displays comparisons of AOX1 expression between Intact and OVX groups, with representative blot images in (C, F). *p<0.05, one way ANOVA, Unpaired t-tests, mean ± SEM, n=6-9/group

